# POLG genotype influences degree of mitochondrial dysfunction in iPSC derived neural progenitors, but not the parent iPSC or derived glia

**DOI:** 10.1101/2023.01.28.526021

**Authors:** Yu Hong, Cecilie Katrin Kristiansen, Anbin Chen, Gonzalo S. Nido, Lena Elise Høyland, Mathias Ziegler, Gareth John Sullivan, Laurence A. Bindoff, Kristina Xiao Liang

**Affiliations:** Department of Clinical Medicine (K1), University of Bergen, Jonas Lies vei 87, P. O. Box 7804, 5021 Bergen, Norway; Neuro-SysMed, Center of Excellence for Clinical Research in Neurological Diseases, Haukeland University Hospital, Jonas Lies vei 87, P. O. Box 7804, 5021 Bergen, Norway; Department of Neurosurgery, Xinhua Hospital Affiliated to Shanghai Jiaotong University School of Medicine, Shanghai, 200092, China; Department of Biomedicine, University of Bergen, Jonas Lies Vei 91, 5009 Bergen, Norway; Department of Molecular Medicine, Institute of Basic Medical Sciences, University of Oslo, P. O. Box 1105, Blindern, 0317 Oslo, Norway; Institute of Immunology, Oslo University Hospital, PO Box 4950, 0424 Oslo, Norway; Hybrid Technology Hub Centre of Excellence, Institute of Basic Medical Sciences, University of Oslo, P. O. Box 1110, Blindern, 0317 Oslo, Norway; Department of Pediatric Research, Oslo University Hospital, P. O. Box 4950, Nydalen, 0424 Oslo, Norway; Department of Neurology, Haukeland University Hospital, Jonas Lies vei 87, P. O. Box 7804, 5021 Bergen, Norway

**Keywords:** POLG, Mitochondrial function, Genotype, Neural stem cells, Neuron

## Abstract

Diseases caused by *POLG* mutations are the most common form of mitochondrial disease and associated with phenotypes of varying severity. Clinical studies have shown that patients with compound heterozygous *POLG* mutations have a lower survival rate than patients with homozygous mutations, but the molecular mechanisms behind this remain unexplored. Using an induced pluripotent stem cell (iPSC) model, we investigate differences between homozygous and compound heterozygous genotypes in different cell types, including patient-specific fibroblasts, iPSCs, and iPSC-derived neural stem cells (NSCs) and astrocytes. We found that compound heterozygous lines exhibited greater impairment of mitochondrial function in NSCs than homozygous NSCs, but not in fibroblasts, iPSCs, or astrocytes. Compared with homozygous NSCs, compound heterozygous NSCs exhibited more severe functional defects, including reduced ATP production, loss of mitochondrial DNA (mtDNA) copy number and complex I expression, disturbance of NAD^+^ metabolism, and higher ROS levels, which further led to cellular senescence and activation of mitophagy. RNA sequencing analysis revealed greater downregulation of mitochondrial and metabolic pathways, including the citric acid cycle and oxidative phosphorylation, in compound heterozygous NSCs. Our iPSC-based disease model can be widely used to understand the genotype-phenotype relationship of affected brain cells in mitochondrial diseases, and further drug discovery applications.

**Graphic Abstract:** 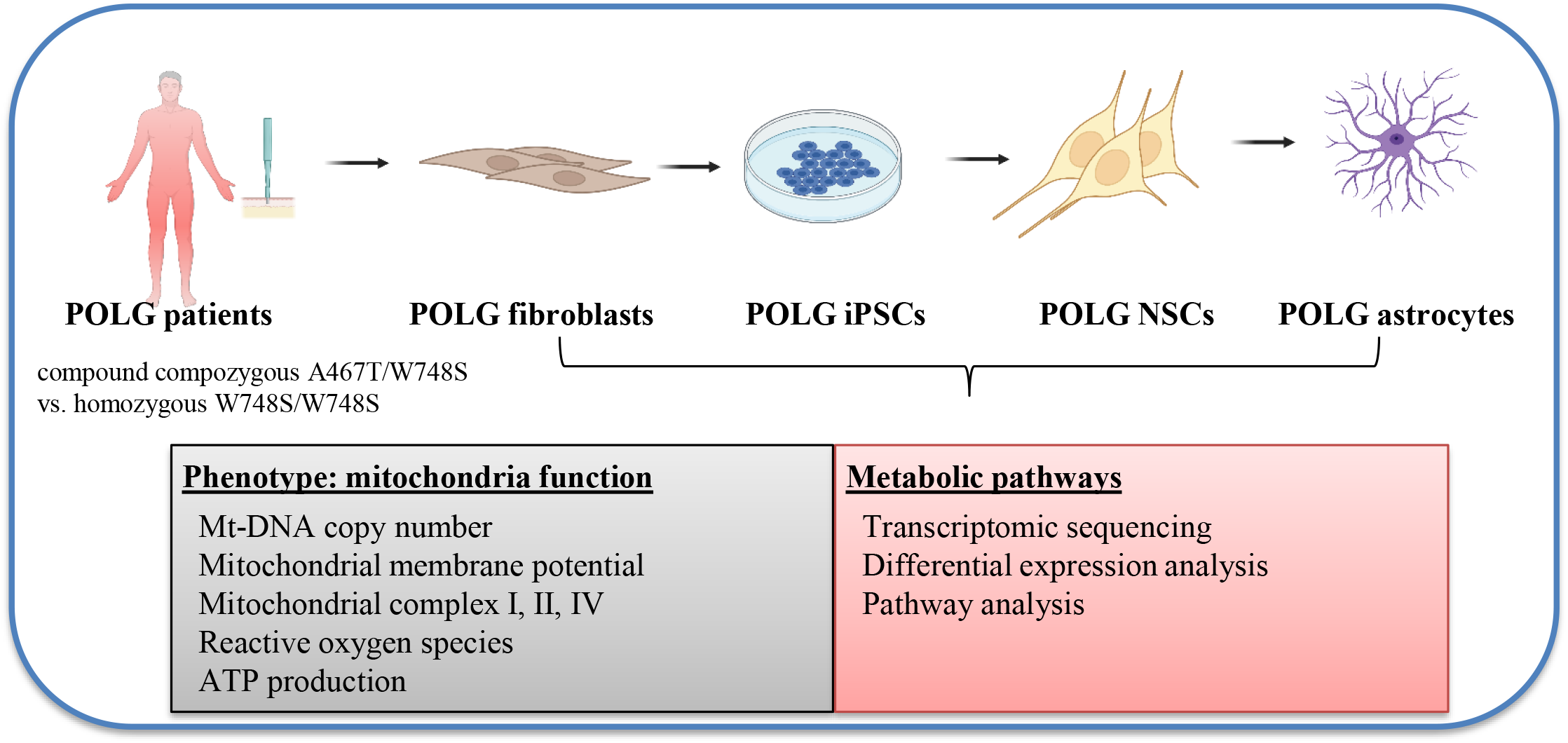

## Introduction

Human mitochondrial DNA (mtDNA) is replicated and repaired by polymerase gamma, Pol γ. Human Pol γ is composed of three subunits encoded by two nuclear genes: POLG gene codes for the 140-kilodalton (kDa) catalytic subunit, p140 and *POLG2* encodes the ~110-kDa homodimeric accessory subunit, p55. The POLG gene encodes the enzyme synthesizing the mitochondrial DNA and correcting mtDNA errors. Mutations in *POLG* gene are the most common causes of inherited mitochondrial diseases and POLG-related disorders comprise a continuum of overlapping phenotypes with onset from infancy to late adulthood. Mitochondrial dysfunction has also been implicated in the pathophysiology of common forms of neurodegeneration such as Parkinson’s disease, therefore, studying how *POLG* mutations affect mitochondrial function and cellular homeostasis is relevant to a variety of diseases.

*POLG* is one of several nuclear genes that are associated with mtDNA depletion or multiple deletion disorder (1), but while mutations in *POLG* can affect the quality or quantity of mtDNA, the type of mtDNA defect depends on the tissue affected in the brain (2, 3). For example, we have shown that severe mtDNA depletion is present in neurons from a very early age, even before morphological changes appear, while mtDNA deletions and point mutations show time-dependent accumulation (2).

To date, more than 300 different diseases associated *POLG* mutations have been identified (1). Among them, c.2243G>C (p.W748S) and c.1399G>A (p.A467T) are two founder mutations defined in multiple populations. Our previous clinical studies suggested that compound heterozygous patients (W748S/A467T) manifested a more severe phenotype than homozygous A467T or W748S (4). However, their genotype-phenotype differences and underlying mechanisms have remained unclear, in part due to the lack of accurate and appropriate models. Human induced pluripotent stem cell (iPSC) technology allows the modeling of human disease in a reproducible and tissue-specific manner without the need for animal models or the use of human material obtained through biopsy or postmortem. We have used this technology to perform molecular studies of the pathogenesis of POLG-related disease.

We previously generated human iPSCs from two POLG patients, one carrying W748S/W748S homozygote (POLG^homo^) and the other harboring compound heterozygous for A467T/W748S (POLG^comp^). Using these lines, we generated neural stem cells (NSCs) and dopaminergic (DA) neurons (5, 6) and showed that they displayed a phenotype that faithfully replicated the molecular and biochemical changes found in postmortem brain tissue of patients (5). Further, we compared these POLG iPSC-derived neuronal cells with healthy controls and found that both POLG^homo^ and POLG^comp^ neuronal cells showed distinct mitochondrial defects, including reduced mitochondrial membrane potential (MMP), ATP depletion, mtDNA depletion, complex I loss, and aberrant NAD^+^ metabolism (5, 6).

In this study, we compared POLG^homo^ and POLG^comp^ cells and analyzed how these genotypes behave in different cell lineages. We investigated mitochondrial function including MMP, respiratory chain complexes, reactive oxygen species (ROS) and mtDNA copy number in fibroblasts, iPSCs and neuronal cells, and astrocytes. In addition, we investigated the transcriptome in cells with phenotypic differences to understand the different impact these genotypes had on other pathways. We found that mitochondrial dysfunction and metabolic pathway dysregulation were more severe in compound heterozygotes, but only in neuronal cells and not in fibroblasts, iPSCs or astrocytes.

## Results

### Compound heterozygous patients have a worse prognosis than those with homozygous mutations

We showed previously that compound heterozygous POLG^comp^ patients had a poorer prognosis than POLG^homo^ patients (7). In order to confirm this, we expanded the sample size by systematically reviewing studies involving patients with both genotypes in Norway, Finland, the UK, Belgium and Sweden (Table S1). We reviewed five published articles (7–11) and analyzed a total of 50 patients, of which 14 were POLG^comp^ patients (7 in Norway, 2 in United Kingdom, 3 in Belgium and 2 in Sweden) and 36 POLG^homo^ patients (13 in Norway, 19 in Finland, 2 in Belgium and 2 in Sweden). Median survival time analysis showed that POLG^comp^ patients (9 years) had significantly (P=0.007) lower survival than POLG^homo^ patients (42 years, Figure S1) confirming our previous observation (7).

### Mitochondrial metabolic pathways are dysregulated in POLG NSCs

IPSCs were reprogrammed from patient fibroblasts using a viral system. We generated NSCs from patient iPSCs via a dual SMAD inhibition approach (Figure 1A), as described previously (5). All iPSCs exhibited typical ESC morphology with sharp edges, however, POLG^comp^ iPSCs exhibited looser packing cells and larger cell size than POLG^homo^ clones (Figure 1B). Flow cytometry analysis showed that all iPSC lines showed that more than 99% of cells stained with the pluripotent stem cell marker OCT4 and SSEA4 (Figure 1C). Immunostaining confirmed that both POLG^homo^ iPSCs (Figure1D) and POLG^comp^ iPSCs (Figure 1E) expressed the specific pluripotent markers OCT4, SOX2 and NANOG. We characterized iPSC derived NSCs using immunostaining for the neural stem cell markers PAX6 and NETSIN, and all NSCs showed positive cells for both markers (Figure 1F). Age-matched iPSC controls and two human embryonic stem cell lines (ESCs): ESC 429 and H1, were used as disease-free controls. To minimize phenotypic diversity caused by intraclonal variation in iPSCs, multiple clones were included in further analyses.

**Figure 1.**
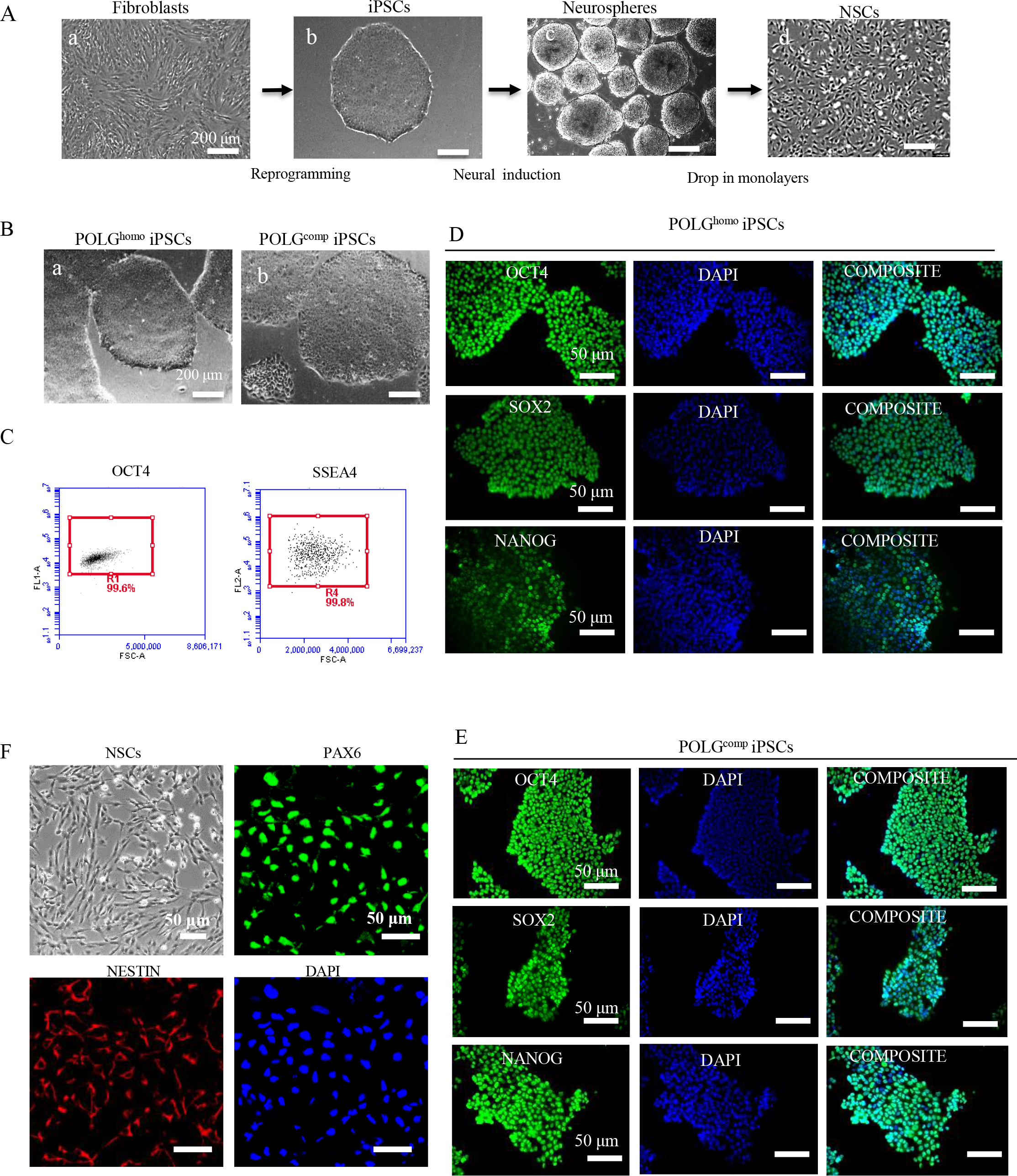
Generation and characterization of iPSCs and NSCs from POLG^homo^ or POLG^comp^ patient. (A). Flow chart of cell reprogramming and differentiation. The fibroblasts (a) were isolated from two POLG patients carrying either POLG^homo^ or POLG^comp^ and reprogrammed into human iPSCs (b). The iPSCs was further differentiated into NSCs (d) via induction of neurospheres (c). (B). Morphology on phase contrast microscopy for iPSC of POLG^homo^ (a) patient and iPSC of POLG^comp^ (b) patient. Scale bars is 200 μm. (C). Flow cytometric quantification of expression level of pluripotent marker OCT4 and SSEA4 in reprogrammed iPSCs. (D, F). Immunofluorescence staining of stem cell markers OCT4 and SOX2 in POLG^homo^ iPSCs (D) and POLG^comp^ iPSCs (F). Scale bar is 50 μm. Nuclei are stained with DAPI (blue).

Next, we compared the transcriptome of iPSCs and their derived NSC’s with controls. Initially, we combined the two genotypes into one patient group and all control lines into one control group for analysis. Sample correlation analysis revealed clear differences between NSC and iPSC/ESC cell types, as well as between the different POLG^homo^ and POLG^comp^ genotypes in each cell type (Figure 2A). Differential expression (DE) analysis of iPSCs and NSCs showed that the number of DE genes in NSCs was much higher than in iPSCs: 1295 upregulated genes and 1263 downregulated genes in patient iPSCs versus control iPSCs, whereas there were 3452 upregulated genes and 3128 downregulated genes in patient NSCs compared to control NSCs (Figure 2B). KEGG module pathway analysis of DE genes revealed that patient NSCs exhibited downregulated mitochondrial oxidative phosphorylation-related pathways, including F- and V-type ATPases, citrate cycle enzymes, NADH dehydrogenase, and cytochrome c oxidation compared with controls (Figure 2C). In contrast, no significantly enriched metabolic pathways were found in patient iPSCs compared to control iPSCs.

**Figure 2.**
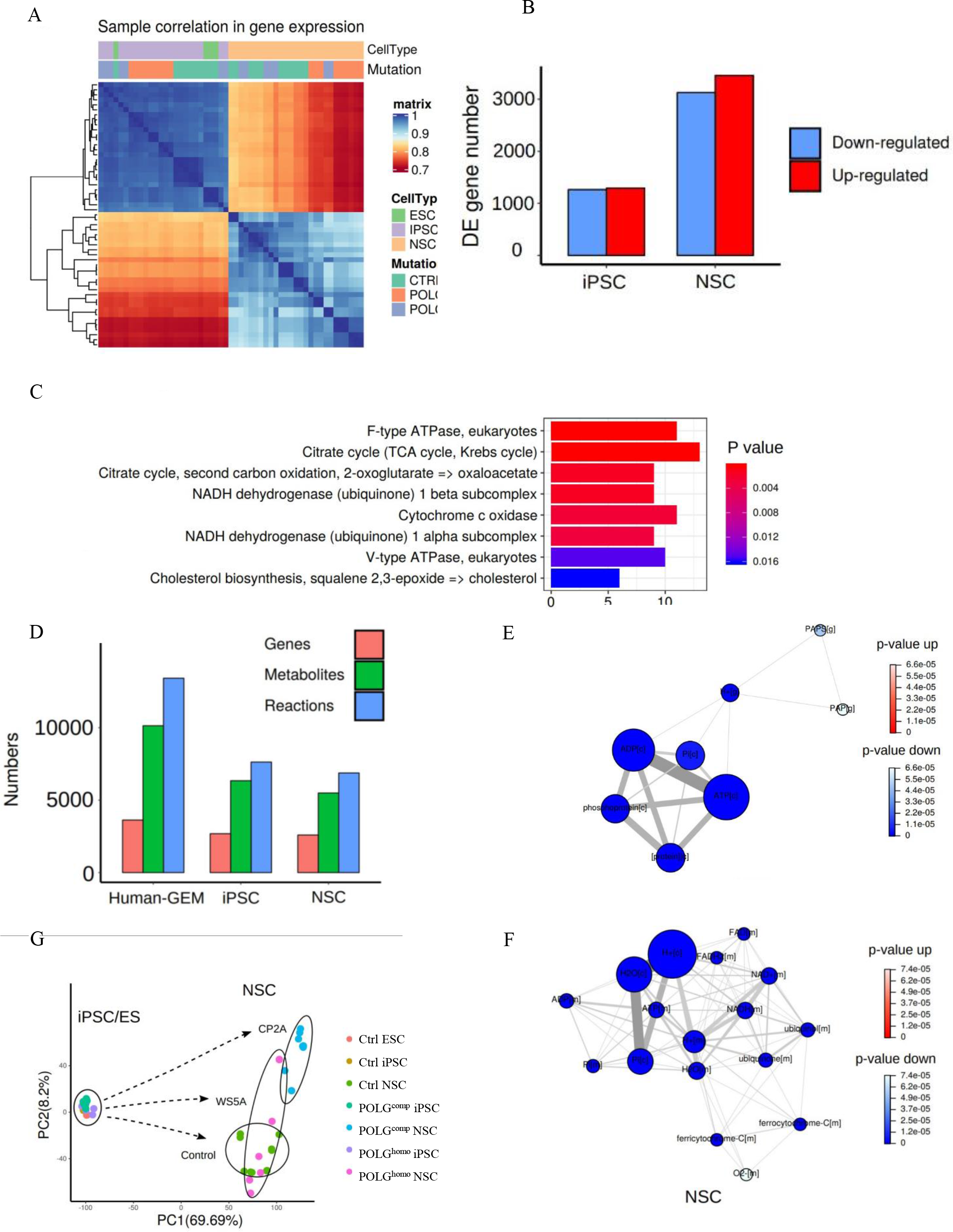
Transcriptomic status and dysregulated metabolic pathways of POLG patients derived iPSCs and NSCs compared to control. (A). Sample correlation in transcriptomic status of POLG patients derived iPSCs and NSCs, and the corresponding controls. (B). Numbers of up- and downregulated DE genes between POLG patients derived iPSCs and NSCs compared to controls, respectively. (C). Significantly enriched KEGG module pathways of downregulated genes in POLG patients derived NSCs compared with controls. (D). Genome scale metabolic models (GEMs) of iPSCs and NSCs, which were constructed based on transcriptomic data of control iPSC/ES and NSCs using a public human genome-scale metabolic model (Human-GEM) as a reference. (E, F). Metabolic reporter analysis of POLG patients derived iPSCs and NSCs based on iPSC and NSC GEMs, respectively. (G). Principal components analysis of transcriptomic matrix for POLG^homo^ and POLG^comp^ patients derived iPSC and NSCs and controls.

To analyze these changes further, we constructed a genome-scale metabolic model of iPSCs and NSCs based on the mRNA expression status of control iPSCs/ES and NSCs using the public human genome-scale metabolic model (Human-GEM). The constructed metabolic model included 7629 reactions, 6342 metabolites, and 2685 genes in iPSCs, and 6877 reactions, 5493 metabolites, and 2594 genes in NSCs (Figure 2D). Metabolic reporter analysis showed that metabolites such as ATP[c], ADP[c], and phosphoprotein[c] were significantly downregulated in POLG patient iPSCs compared to control iPSCs (Figure 2E). When comparing patient and control NSCs, we found downregulation of metabolic proteins in patient NSCs, including H2O[c], H+[c], H2O[c], NAD+[m], NADH[m], FAD+[m], and FADH2 [m] (Figure 2F).

We next explored differences in gene expression profile in iPSCs and NSCs carrying POLG^homo^ and POLG^comp^ mutations. We performed principal component analysis (PCA) for POLG^homo^, POLG^comp^ and control lines and found that POLG^homo^, POLG^comp^ and control iPSCs had similar gene expression profiles (Figure 2G). In NSCs, there was greater heterogeneity among the three groups (Figure 2G). More notably, more genes were dysregulated in between POLG^comp^ NSCs and controls than between POLG^homo^ NSCs and controls (Figure 2G).

These data suggest that the metabolic dysregulation induced by *POLG* mutations only manifests as cells mature toward neuronal lineage

### Compound heterozygous and homozygous iPSCs show similar mitochondrial function and transcriptomic profiling

We profiled pluripotency using stem cell marker genes and protein expression of pluripotent markers. QPCR analysis showed similar expression of *LIN28A*, *POU5F1* and *NANOG* in POLG^homo^ and POLG^comp^ iPSCs (Figure 3A). Flow cytometry also detected similar expression levels of all pluripotency markers in both mutant iPSCs, including the pluripotent surface markers SSEA4, TRA-1-60 and TRA-1-81 (Figure 3B) and the transcription factors OCT4, NANOG (Figure 3C).

**Figure 3.**
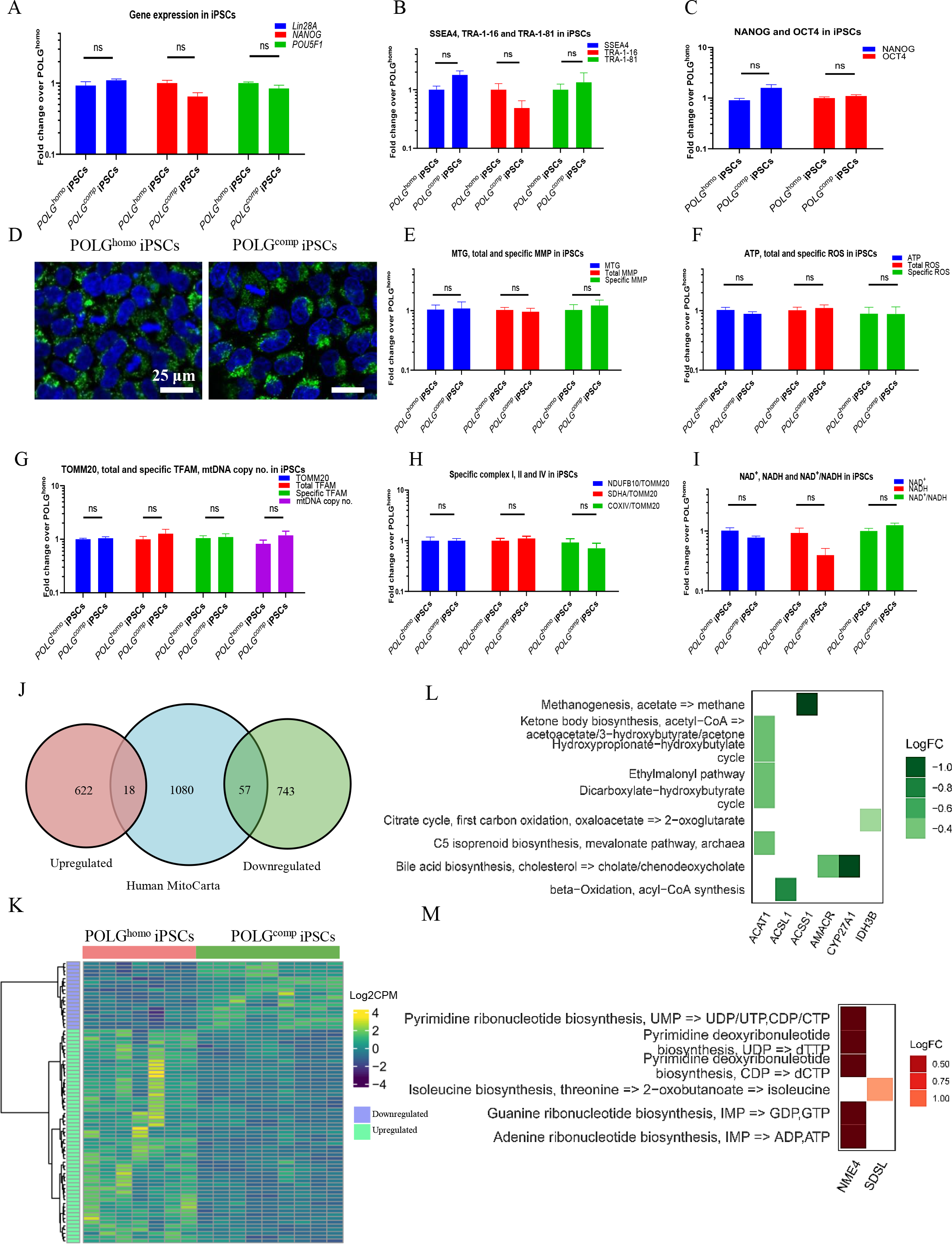
Measurement of mitochondrial function, mtDNA alteration and NAD^+^ metabolism in POLG^homo^ and POLG^comp^ iPSCs. (A). QPCR quantification of gene expression for *LIN28A*, *NANOG*, and *POU5F1* for POLG^homo^ and POLG^comp^ patients iPSCs. The fold change is calculated by normalizing all the values to the mean value of POLG^homo^ iPSCs. (B, C). Flow cytometric analysis for pluripotent surface markers SSEA4, TRA-1-60, andTRA-1-81(B), as well as transcription factors NANOG and OCT4 (C) in POLG^comp^ iPSCs compared with POLG^homo^ iPSCs. The fold change is calculated by normalizing all the values to the mean value of POLG^comp^ iPSCs. (D). Confocal images of mitochondrial morphology for POLG^homo^ and POLG^comp^ iPSC lines with MTG staining. Scale bars is 25 μm. Nuclei are stained with DAPI (blue). (E). Flow cytometric analysis of iPSCs for mitochondrial volume (MTG), total MMP (TMRE) and specific MMP (TMRE/MTG) levels. The fold change is calculated by normalizing all the values to the mean value of POLG^homo^ iPSCs. (F). Measurement of intracellular ATP production from luminescent assay and flow cytometric analysis for total intracellular ROS level (DCFDA), and the specific ROS level (DCFDA/MTDR) for POLG^homo^ and POLG^comp^ iPSCs. The fold change is calculated by normalizing all the values to the mean value of POLG^homo^ iPSCs. (G). Flow cytometric analysis for TOMM20, total TFAM and specific TFAM (total TFAM/TOMM20) levels, and relative mtDNA copy number analyzed by QPCR for POLG^homo^ and POLG^comp^ iPSCs. The fold change is calculated by normalizing all the values to the mean value of POLG^homo^ iPSCs. (H). Flow cytometric measurements of mitochondrial complex I, II and IV protein expression at total ad specific level (total complex I, II, IV level/TOMM20). The fold change is calculated by normalizing all the values to the mean value of POLG^homo^ iPSCs. (I). LC-MS-based metabolomics for quantitative measurement of total NAD^+^, total NADH level and NAD^+^/NADH ratio in POLG^homo^ and POLG^comp^ iPSCs. The fold change is calculated by normalizing all the values to the mean value of POLG^homo^ iPSCs. (J). DE analysis between POLG^homo^ and POLG^comp^ iPSCs using the human MitoCarta 2.0 database was used as a reference of mitochondrial function related genes. (K). Heatmap of DE expressed mitochondrial function related genes in POLG^homo^ and POLG^comp^ iPSCs. (L-M). KEGG metabolic pathway module classification of up-(L) and downregulated (M) mitochondrial functional related DE genes POLG^homo^ and POLG^comp^ iPSCs. Y-axis represents pathway modules and X-axis represents mitochondrial related DE genes belong to each module. However, none of these pathway modules were significantly enriched.

Using MitoTracker Green (MTG) staining and fluorescence confocal microscopy, we determined POLG^homo^ and POLG^comp^ iPSCs had similar morphology (Figure 3D). Next, we used our previously established flow cytometry method (12) to measure mitochondrial function. We quantified the levels of MTG and tetramethylrhodamine ethyl ester (TMRE) to measure mitochondrial mass and MMP and found that mass, measured by MTG, total MMP, measured by TMRE and specific MMP calculated by TMRE/MTG were similar in POLG^homo^ and POLG^comp^ iPSCs (Figure 3E). ATP production per cell using a live cell luminescence assay was also similar (Figure 3F) as was ROS production measured using dual staining with 2′, 7′-dichlorodihydrofluorescein diacetate (DCFDA) and MitoTracker Deep Red (MTDR) (Figure 3F). To ensure that this reflected the mitochondrial mass in each cell type, we divided total ROS by a measure of MTDR to give specific ROS and again found no difference between these two mutant iPSCs (Figure 3F).

We then investigated mitochondrial respiratory chain complex expression in these two mutant iPSCs. We used the complex I subunit NDUFB10 antibody and co-stained with TOMM20 to correlate complex levels with mitochondrial mass and found similar levels of complex I in POLG^homo^ and POLG^comp^ iPSCs (Figure 3H). Complex II levels using anti-SDHA and complex IV levels measured by anti-COXIV were also similar in both mutant iPSCs (Figure 3H). Lastly, we examined the redox state by measuring NAD^+^, NADH levels and NAD^+^/NADH ratio using liquid chromatography mass spectrometry (LC-MS) and again, found no difference between POLG^homo^ and POLG^comp^ iPSCs (Figure 3I).

Next, we qualitatively and quantitatively assessed mtDNA using flow cytometric analysis of TFAM and qPCR (5). Flow cytometry measurements of TFAM and TOMM20 showed no differences in TOMM20, total TFAM levels, and specific TFAM levels (TFAM/TOMM20) in POLG^homo^ and POLG^comp^ iPSCs (Figure 3G). QPCR measurement of the ND1/APP ratio also revealed no difference in mtDNA copy number in these two mutant iPSCs (Figure 3G).

To explore differences in gene expression, we performed RNA sequencing analysis and compared the expression profiles of POLG^homo^ and POLG^comp^ iPSCs. Although PCA showed similarities in both mutant iPSCs (Figure 1), we identified 1440 DE genes in POLG^comp^ compared to POLG^homo^ iPSCs: 640 genes were upregulated, and 800 genes were downregulated (Figure 3J). Applying these data to the human MitoCarta 2.0 database (13) identified 75 DE genes with purported mitochondrial function, of which 18 were upregulated and 57 were downregulated (Figure 3J, K, Table S2–S3). Subsequent KEGG module analysis showed that the *ACAT1* gene involved in ketone body synthesis and the *IDH3B* gene encoding isocitrate dehydrogenase isoenzyme were significantly downregulated (Figure 3L) while the *SDSL* and *NME4* genes involved in isoleucine and ribonucleotide biosynthesis were significantly upregulated (Figure 3M). Despite some changes in modular pathways, metabolic pathway analysis showed no enrichment when comparing POLG^comp^ iPSCs to POLG^homo^ iPSCs.

These data suggest that compound heterozygous and homozygous POLG iPSCs have similar mitochondrial function and metabolic pathway profiles.

### Mitochondrial function impairment is greater in compound heterozygous NSC than homozygous NSCs

We first examined the purity of the iPSC derived NSC lineage using flow cytometry and double staining with the neural progenitor markers PAX6 and NESTIN. This showed that 99.9% of cells were positive for both PAX6 and NESTIN (Figure 4A). Quantification of protein expression revealed significantly elevated levels of PAX6, but lower levels of NESTIN in POLG^comp^ NSCs compared to POLG^homo^ NSCs (Figure 4B). We assessed mitochondrial structure by transmission electron microscopy and found that both mutant NSCs had similar mitochondrial appearance (Figure 4C). Using the TMRE/MTG double staining method described above, we found that mitochondrial mass (MTG), total MMPs (total TMRE) were similar in these two mutant NSC, but the specific MMP levels, i.e., MMPs per mitochondrial mass (total TMRE/MTG) was lower in POLG^comp^ NSCs compared to POLG^homo^ NSCs (Figure 4D). Direct measurement of intracellular ATP production showed significantly reduced levels in POLG^comp^ NSCs (Figure 4E). The total and specific levels of intercellular ROS and the total mitochondrial ROS level appeared to be similar, however, POLG^comp^ NSCs had lower specific mitochondrial ROS level compared to POLG^homo^ NSCs (Figure 4E).

**Figure 4.**
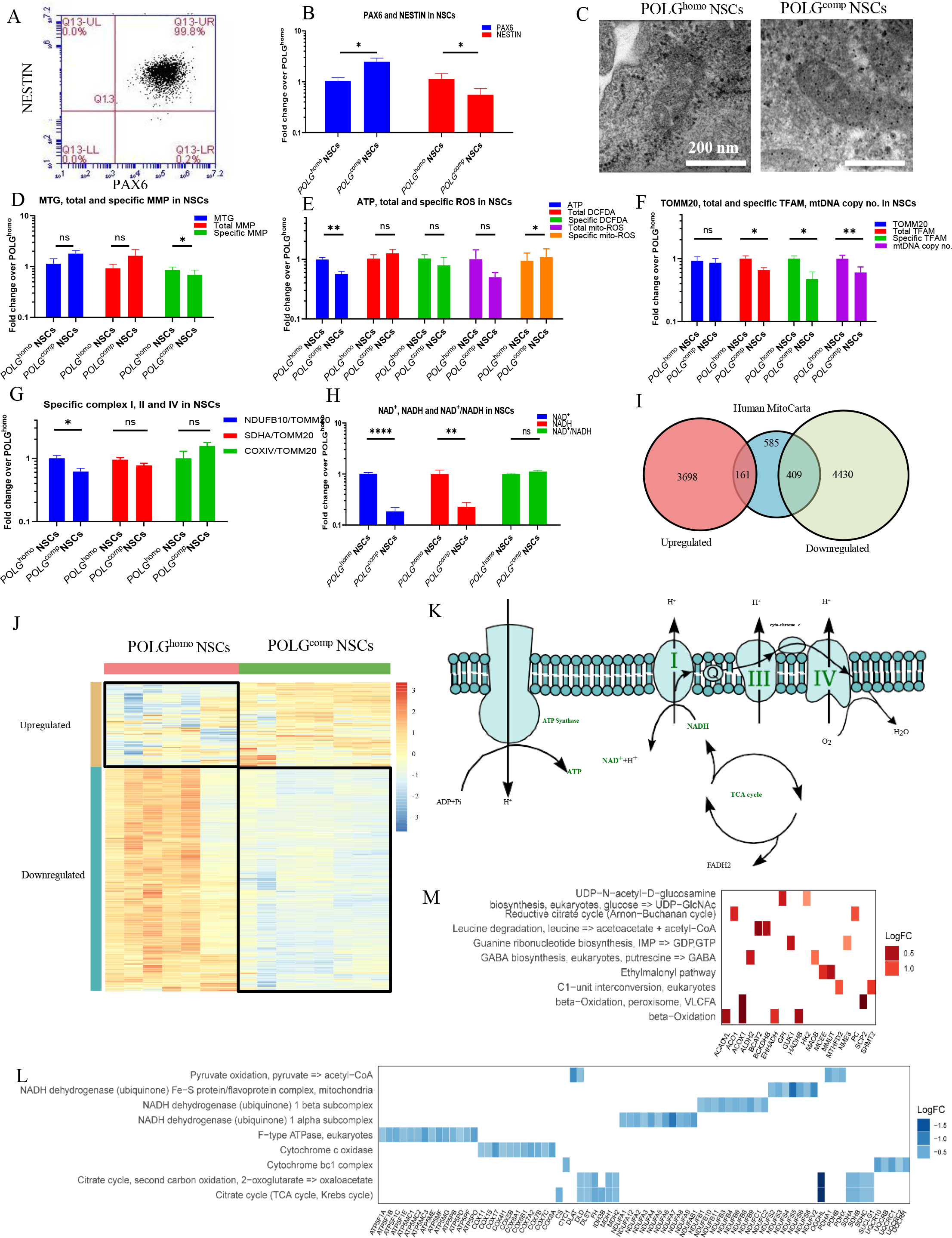
Measurement of mitochondrial function, mtDNA alteration and NAD^+^ metabolism in POLG^homo^ and POLG^comp^ NSCs. (A). Flow cytometric analysis for double staining of the NESTIN and PAX6 in iPSC derived NSCs. (B). Flow cytometric analysis for the expression of NESTIN and PAX6 in POLG^comp^ NSCs compared with POLG^homo^ NSCs. The fold change is calculated by normalizing all the values to the mean value of POLG^comp^ NSCs. (C). Representative transmission electron microscopy images of mitochondrial structures in POLG^homo^ and POLG^comp^ NSCs Scale bar is 200 nm. (D). Flow cytometric analysis of iPSCs for mitochondrial volume (MTG), total MMP (TMRE) and specific MMP (TMRE/MTG) levels. The fold change is calculated by normalizing all the values to the mean value of POLG^comp^ NSCs. (E). Measurement of intracellular ATP production from luminescent assay and flow cytometric analysis for total intracellular ROS level (DCFDA), and the specific ROS level (DCFDA/MTDR) for POLG^homo^ and POLG^comp^ NSCs. The fold change is calculated by normalizing all the values to the mean value of POLG^comp^ NSCs. (F). Flow cytometric analysis for TOMM20, total TFAM and specific TFAM (total TFAM/TOMM20), and relative mtDNA copy number analyzed by qPCR for POLG^homo^ and POLG^comp^ NSCs. The fold change is calculated by normalizing all the values to the mean value of POLG^comp^ NSCs. (G). Flow cytometric measurements of mitochondrial complex I, II and IV protein expression at specific level (total complex I, II, IV level/TOMM20) in POLG^homo^ and POLG^comp^ NSCs. The fold change is calculated by normalizing all the values to the mean value of POLG^comp^ NSCs. (H). LC-MS-based metabolomics for quantitative measurement of total NAD^+^, total NADH level and NAD^+^/NADH ratio in POLG^homo^ and POLG^comp^ NSCs. The fold change is calculated by normalizing all the values to the mean value of POLG^homo^ NSCs. (I). DE analysis between POLG^homo^ and POLG^comp^ NSCs using the human MitoCarta 2.0 database was used as a reference of mitochondrial function related genes. (J). Heatmap of DE expressed mitochondrial function related genes in NSCs between POLG^homo^ and POLG^comp^ lines. (K). Illustration of downregulated mitochondrial functional related pathway modules and metabolites in POLG^comp^ compared with POLG^homo^ NSCs. Involved pathways and metabolites are in green. (L). KEGG metabolic pathway module classification of downregulated DE expressed mitochondrial functional genes in NSCs in POLG^comp^ compared with POLG^homo^ NSCs. (M). KEGG metabolic pathway module classification of upregulated DE expressed mitochondrial functional genes in POLG^comp^ NSCs compared with POLG^homo^ NSCs. Y-axis represents pathway modules and X-axis represents mitochondrial related DE genes belong to each module. (L). KEGG metabolic pathway module classification of downregulated DE expressed mitochondrial functional genes in POLG^comp^ NSCs compared with POLG^homo^ NSCs. Asterisk represents significant enriched pathways. Y-axis represents pathway modules and X-axis represents mitochondrial related DE genes belong to each module.

Analysis of mtDNA as described above showed similar expression of TOMM20 and total TFAM, but decreased levels of specific TFAM, suggesting lower mtDNA amount, which was confirmed by qPCR that showed lower mtDNA copy number in POLG^comp^ (ND1/APP) (Figure 4F). Flow cytometry and TOMM2O co-staining revealed greater loss of specific complex I level in POLG^comp^ NSCs than POLG^homo^ NSCs, while specific levels of complex II and IV were the same (Figure 4G). Furthermore, LC-MS measurements showed that NAD^+^ level and NADH level were lower in POLG^comp^ NSCs compared to POLG^homo^ NSCs, while the ratio of NAD^+^/NADH levels remained unchanged (Figure 4H).

These data suggest that the presence of compound heterozygous *POLG* mutations in NSCs results in greater mitochondrial functional dysfunction than occurs with homozygous mutations.

### Compound heterozygous NSCs showed greater transcriptome defects than homozygous NSCs

Next, we asked whether different NSC genotypes displayed distinct transcriptomic signatures: PCA analysis revealed a clear difference in PCA profiles between POLG^comp^ and POLG^homo^ NSCs (Figure 2G). For comparison, we used 6 POLG^comp^ samples with 2 individual clones each and 3 replicates each, and 9 POLG^homo^ samples with 3 individual clones, 3 replicates each. There were 8698 DE genes in POLG^comp^ NSCs, of which 4839 were downregulated and 3859 were up-regulated (Figure 4I). Enrichment for mitochondrial function-related genes detected 161 upregulated and 409 downregulated genes (Figure 4I, Tables S4–S5). This predominance of downregulated DE genes (Figure 4J) in POLG^comp^ NSCs comprised genes in key mitochondrial functional modules involved in electron transport and energy production processes (Figure 4K, Tables S4–S5). Notably, key regulatory genes involved in pyruvate oxidation (pyruvate => acetyl-CoA) including genes encoding different components of the pyruvate dehydrogenase complex, such as *E1 (PDHA1, PDHB)*, *E2 (DLAT)*, *E3 (DLD)* and *PDHX* (Figure 4L). Additional downregulated genes included ones involved in encoding oxoglutarate dehydrogenase complex (*OGDHL*, *DLD*, *DLST*), succinate-CoA ligase genes (*SUCLG1*), succinate dehydrogenases (SDHA, SDHB, SDHC), and fumarate (FH) and malate dehydrogenases (*MDH1*, *MDH2*) (Figure 4L).

Consistent with our functional investigations, we found downregulation of the genes encoding respiratory chain complex I proteins including *NDUFA1-A8*, β-subcomplex-related genes *NDUFB1, NDUFB3-B4, NDUFB6, NDUFB8-B11, NDUFC1* and the mitochondrial Fe-S protein-related genes *NDUFS2-S6*, *NDUFS8*, and *NDUFV2* (Figure 4L). Multiple genes encoding respiratory chain complex V proteins including catalytic core-related proteins (including *ATP5F1A*, *ATP5F1B*, *ATP5F1C*, *and ATP5F1E*) and proton channel subunit-related proteins (including *ATP5PB*, *ATP5PD*, *ATP5PF*, *ATP5PO*) and those involved in *F1/Fo ATPase* fundtion (including *ATP5MC1-C3, ATP5ME*, *ATP5MF and ATP5MG*) (Figure 4L) were also downregulated. Genes encoding complex III proteins (cytochrome bc1*: CYC1*, *UQCRB*, *UQCRC1* and *UQCRC2*, *UQCR10*, *UQCRH, etc.*) and complex IV proteins (cytochrome c oxidase: *COX6B1*, *COX7A2*, *COX8A, etc.*) were also downregulated (Figure 4L). Pathway enrichment analysis of KEGG modules revealed that downregulated genes in POLG^comp^ NSCs were mainly distributed in β-oxidation (including *ACADVL*, *ACOX1*, *EHHADH*, *HADHB*), proline biosynthesis (including *ALDH18A1* and *PYCR1*) and isoleucine biosynthesis (including *BCAT2* and *SDSL*). (Figure 4L). Although there were upregulated genes in POLG^comp^ NSCs relative to POLG^homo^ NSCs, these were not significantly different (Figure 4L).

These findings suggest that compound heterozygous mutations in POLG patients lead to more severe dysregulation of metabolic pathways in NSCs.

### Greater mitochondrial defects in compound heterozygous NSCs led to cellular senescence and mitophagy activation

Since our findings suggested greater mitochondrial dysfunction in compound heterozygous NSCs, we asked what the cellular consequences of these changes were. Given changes in NAD^+^ metabolism and complex I and the known link between mitochondrial dysfunction and increased ROS production and senescence, we investigated cellular senescence and mitophagy in our NSCs. To examine cellular senescence, we performed β-galactosidase staining and flow cytometry to measure the cellular senescence marker p16INK4. We observed a significant increase in β-galactosidase activity in compound heterozygous versus homozygous NSCs (Figure 5A, B). This finding was further confirmed by p16INK4 expression measured by flow cytometry (Figure 5B).

**Figure 5.**
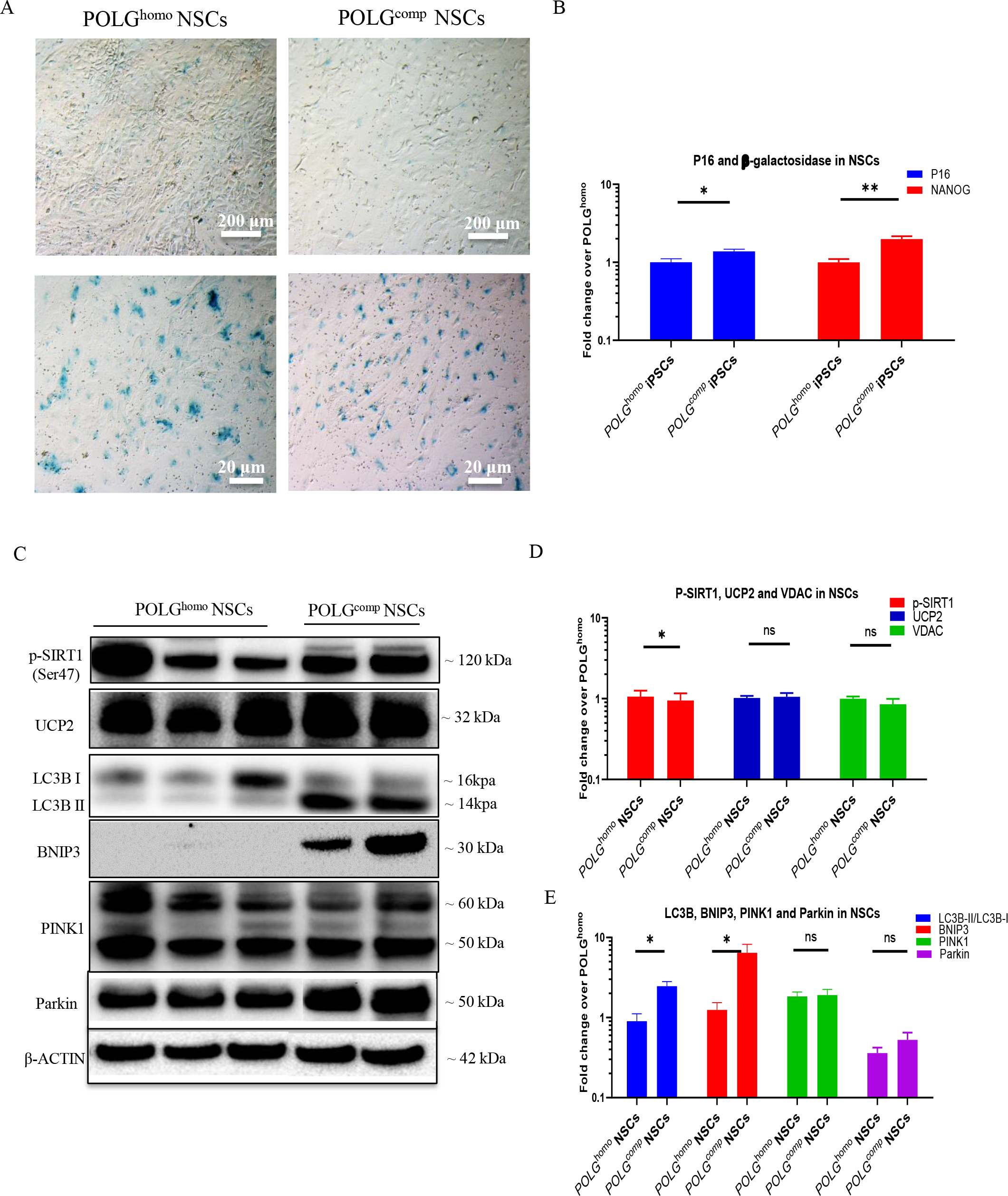
POLG^comp^ NSCs exhibited senescence phenotype and mitophagy activation compared to POLG^homo^ NSCs. (A, B). Representative images of senescence β-galactosidase staining (A) and quantification (B) by calculating the percentage of positively stained cells by division of the negative cells from A in POLG^comp^ NSCs compared to POLG^homo^ NSCs. Scale bars is 20 μm or 200 μm. The fold change in B is calculated by normalizing all the values to the mean value of POLG^homo^ NSCs. (C-E). Representative images (C) and quantification (D, E) for western blotting with Phospho-SirT1 (Ser47), UCP2, VDAC, LC3B, BNIP3, PINK1, Parkin and β-ACTIN. Three independent experiments are included. The fold change in D and E is calculated by normalizing all the values to the mean value of POLG^homo^ NSCs.

Next, we investigated the molecular pathways that initiate cellular senescence. Sirtuin 1 (SIRT1) is an NAD^+^-dependent deacetylase that plays a key role in many cellular processes including energy metabolism, stress response and aging (14). SIRT1 also regulates the expression of the mitochondrial uncoupling protein 2 (*UCP2*) gene, overexpression of which can reduce MMPs and induce senescence (14). We investigated the expression of phosphorylated SIRT1 (Ser47) (p-SIRT1) and UCP2 using western blotting and found that phosphorylated SIRT1 was reduced in POLG^comp^ NSCs compared to POLG^homo^ NSCs, altough UCP2 expression remained unchanged (Figure 5C, D).

Mitophagy, or autophagy, is required for both promotion and inhibition of senescence and senescence-related phenotypes (15). To investigate whether mitophagy was altered, we investigated the mitophagy-related proteins LC3B, BNIP3, PINK1 and Parkin using western blotting. This revealed elevated level of LC3B-II and elevated LCBII/LC3BI ratio in POLG^comp^ NSCs, indicating autophagy activation (Figure 5C, E). Accumulation of BNIP3 was also observed in POLG^comp^ NSCs, but not in POLG^homo^ cells (Figure 5C, E). The expression of PINK1 and Parkin was similar in both mutant NSCs (Figure 5C, E).

These findings suggest that the more severe mitochondrial damage in heterozygous POLG variants results in increased cellular senescence through downregulation of p-SIRT1 and activation of mitophagy or autophagy regulated by BNIP3.

### The effect on mitochondrial function in astrocytes is similar in compound heterozygous and homozygous

Next, we tested the effect of different genotypes in astrocytes. We differentiated astrocytes from NSCs using a previously described protocol (16). Cell morphology was similar in compound heterozygous and homozygous astrocytes (Figure 6A) and confocal images showed positive staining for astrocyte lineage markers GFAP and S100β and astrocyte functional markers EAAT1 and glutamine synthesis (GS) (Figure 6A). Differentiated astrocytes also stained negative for the neural marker dcx (DCX) (Figure S3), indicating no contamination of neural lineages. Flow cytometric analysis confirmed that astrocyte lineages were > 98% positive for lineage markers GFAP and CD44 and functional markers EAAT1 and GS (Figure 6B). Quantification of the protein levels of these astrocyte markers showed similar levels of GFAP, CD44, S100β, EAAT1, and GS in two mutant astrocytes (Figure 6C).

**Figure 6.**
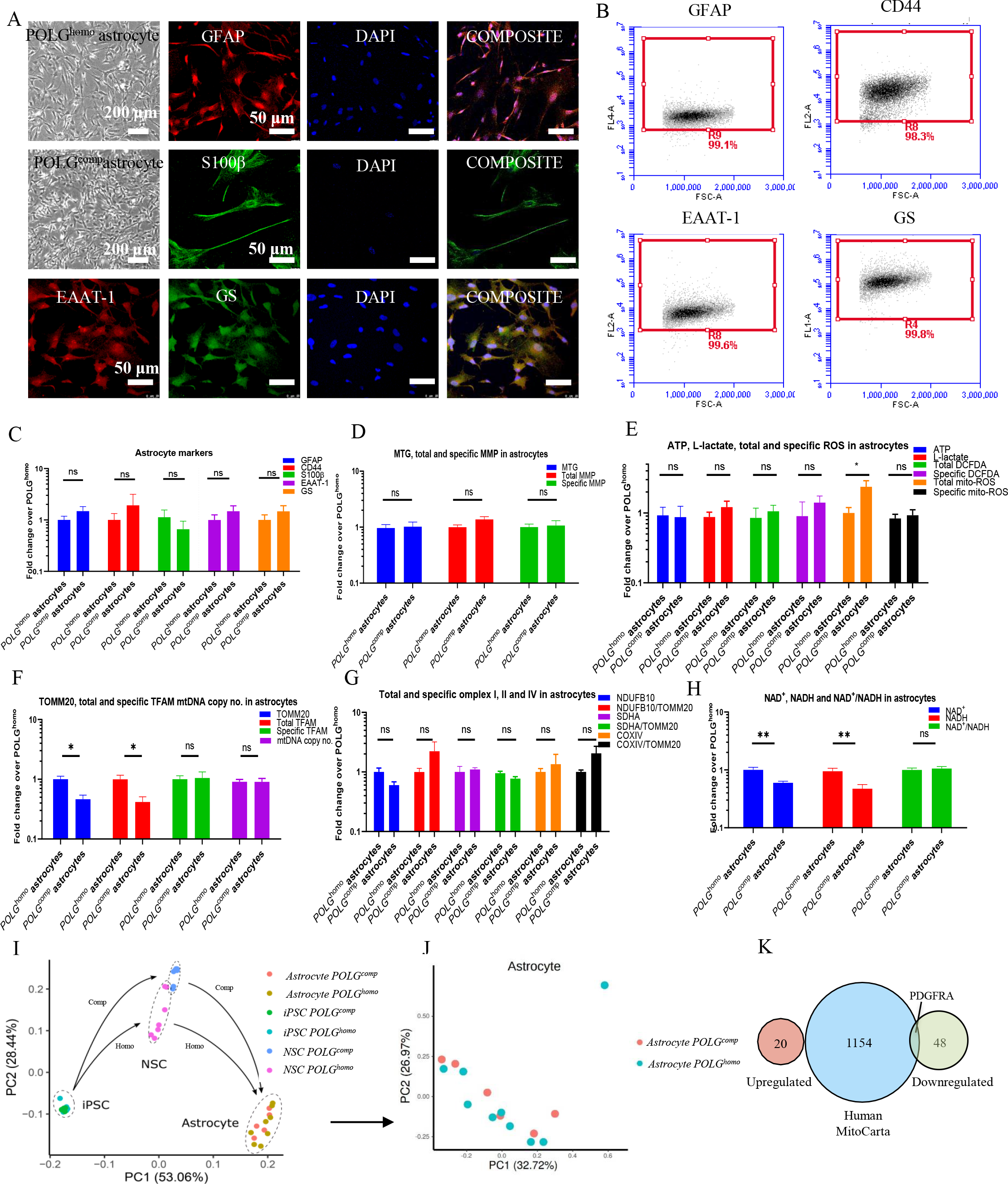
Measurement of mitochondrial function, mtDNA alteration and NAD^+^ metabolism in POLG^homo^ and POLG^comp^ astrocytes. (A). Representative phase-contrast images of the iPSC-derived astrocytes generated from in POLG^comp^ and POLG^homo^ astrocytes and confocal images of astrocyte lineage markers GFAP and S100β, and functional markers EAAT-1 and GS. Scale bar is 200 μm or 50 μm. Nuclei are stained with DAPI (blue). (B). Flow cytometric analysis for GFAP, CD44, EAAT-1 and GS in astrocytes. (C). Flow cytometric analysis for the expression of GFAP, CD44, S100β, EAAT-1 and GS in POLG^comp^ astrocytes compared with POLG^homo^ astrocytes. The fold change is calculated by normalizing all the values to the mean value of POLG^homo^ astrocytes. (D). Flow cytometric analysis for mitochondrial volume (MTG), total MMP (TMRE) and specific MMP (TMRE/MTG) levels in POLG^comp^ astrocytes compared with POLG^homo^ astrocytes. The fold change is calculated by normalizing all the values to the mean value of POLG^homo^ astrocytes. (E). Measurement of intracellular ATP production from luminescent assay and flow cytometric analysis for total intracellular ROS level (DCFDA), and the specific ROS level (DCFDA/MTDR) for POLG^homo^ and POLG^comp^ astrocytes. The fold change is calculated by normalizing all the values to the mean value of POLG^homo^ astrocytes. (F). Flow cytometric analysis for TOMM20, total TFAM and specific TFAM (total TFAM/TOMM20), and relative mtDNA copy number analyzed by qPCR for POLG^homo^ and POLG^comp^ astrocytes. The fold change is calculated by normalizing all the values to the mean value of POLG^homo^ astrocytes. (G). Flow cytometric measurements of mitochondrial complex I, II and IV protein expression at total and specific level (total complex I, II, IV level/TOMM20) in POLG^homo^ and POLG^comp^ astrocytes. The fold change is calculated by normalizing all the values to the mean value of POLG^homo^ astrocytes. (H). LC-MS-based metabolomics for quantitative measurement of total NAD^+^, total NADH level and NAD^+^/NADH ratio in POLG^homo^ and POLG^comp^ astrocytes. The fold change is calculated by normalizing all the values to the mean value of POLG^homo^ astrocytes. (I). PCA analysis of POLG^comp^ and POLG^homo^ iPSCs, NSCs and astrocytes. (J). PCA analysis of POLG^comp^ and POLG^homo^ astrocytes. (K). DE analysis between POLG^homo^ and POLG^comp^ astrocytes using the human MitoCarta 2.0 database was used as a reference of mitochondrial function related genes.

Using the same experimental approach as described above, we found that POLG^comp^ and POLG^homo^ astrocytes exhibited similar mitochondrial mass (MTG) and total and specific levels of MMP (Figure 6D). Although lower levels of total mitochondrial ROS were detected in POLG^comp^ astrocytes versus POLG^homo^ astrocytes, no difference in levels of ATP production, lactate production, total and specific intracellular ROS, specific mitochondrial ROS were detected between the two mutant astrocytes (Figure 6E). We found lower expression of TOMM20 and total TFAM in compound heterozygous cells (Figure 6F), but there was no change in specific TFAM level (TFAM normalized to mitochondrial content), and qPCR measurement showed no difference in mtDNA copy number between genotypes (Figure 6F). Measurements of the complexes I-IV at both total and specific levels showed no difference between the two mutant astrocytes (Figure 6G). Measurements of NAD biosynthesis showed significantly reduced levels of NAD^+^ and NADH in POLG^comp^ astrocytes versus POLG^homo^ astrocytes, but the ratio of NAD^+/^NADH was similar (Figure 6H).

We then investigated the transcriptional signature of POLG^comp^ and POLG^homo^ astrocytes. We found in PCA analysis that although NSCs from both genotypes showed greater heterogeneity, astrocytes differentiated from the two groups of NSCs did not differ significantly in astrocyte RNA expression (Figure 7I, J). For DE analysis, we found that the expression of only 69 genes was significantly altered in when comparing POLG^comp^ to POLG^homo^ astrocytes, including 20 upregulated genes and 49 downregulated genes (Figure 7K). Notably, only one gene, *PDGFRA*, belonging to the MitoCarta mitochondrial gene database, was downregulated in POLG^comp^ astrocytes than in POLG^homo^ astrocytes (Figure 7K).

**Figure 7.**
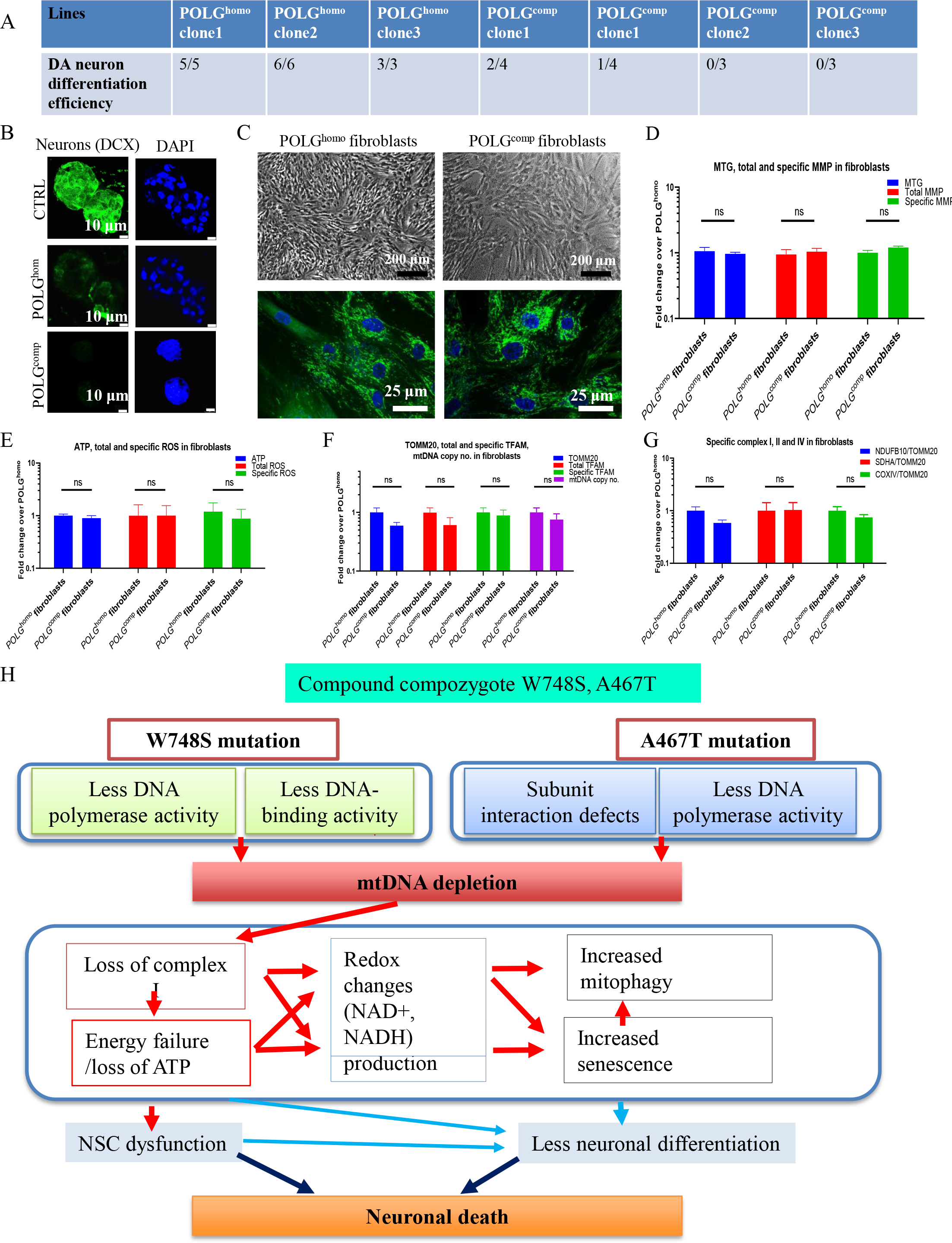
DA neural differentiation efficiency of POLG^homo^ and POLG^comp^ cells and measurement of mitochondrial function, mtDNA alteration in POLG^homo^ and POLG^comp^ fibroblasts. (A). DA neuron differentiation efficiency in induvial clones from POLG^homo^ and POLG^comp^ iPSCs. (B). Representative confocal microscopy images of DCX staining of neurospheres made from NSCs, astrocytes and oligodendrocytes derived from POLG^homo^, POLG^comp^ and control iPSCs. (C). Representative phase-contrast and confocal microscopy images with MTG staining in POLG^homo^ and POLG^comp^ fibroblasts. Scale bar is 200 μm or 25 μm. Nuclei are stained with DAPI (blue). (D). Flow cytometric analysis for MTG, total MMP (TMRE) and specific MMP (TMRE/MTG) in POLG^homo^ and POLG^comp^ fibroblasts. The fold change is calculated by normalizing all the values to the mean value of POLG^homo^ fibroblasts. (E). Measurement of intracellular ATP production from luminescent assay and flow cytometric analysis for total intracellular ROS level (DCFDA), and the specific ROS level (DCFDA/MTDR) for POLG^homo^ and POLG^comp^ fibroblasts. The fold change is calculated by normalizing all the values to the mean value of POLG^homo^ fibroblasts. (F). Flow cytometric analysis for TOMM20, total TFAM and specific TFAM (total TFAM/TOMM20), and relative mtDNA copy number analyzed by qPCR for POLG^homo^ and POLG^comp^ fibroblasts. The fold change is calculated by normalizing all the values to the mean value of POLG^homo^ fibroblasts. (G). Flow cytometric measurements of mitochondrial complex I, II and IV protein expression at total and specific level (total complex I, II, IV level/TOMM20) in POLG^homo^ and POLG^comp^ fibroblasts. The fold change is calculated by normalizing all the values to the mean value of POLG^homo^ fibroblasts. (H). Summary of the possible disease mechanisms in neuronal death with the patient carrying *POLG* compound heterozygous mutation.

These studies demonstrate that the genotype-defined mitochondrial and transcriptional differences seen in NSCs are lost in the astrocyte lineage, even though astrocytes are derived from NSCs.

### Compound heterozygous POLG NSCs have a lower potential for differentiation to DA neurons than homozygous NSCs

Our previous clinical study showed that dopaminergic (DA) neurons in the substantia nigra are among the most severely affected cell types in POLG patients (17). We also reported previously that NSC-derived DA neurons exhibit defective mitochondrial function and mtDNA depletion with a trend suggesting a greater defect in POLG^comp^ than in POLG^homo^ neurons (6). In this current study, we asked whether differences in mitochondrial damage between compound heterozygous and homozygous NSCs affected the differentiation efficiency towards the commitment of DA neurons. To do this, we used 3 separate POLG^homo^ NSCs clones and 4 separate POLG^comp^ NSCs clones and performed DA differentiation multiple times. POLG^comp^ NSCs were consistently less efficient at differentiating neurons than POLG^homo^ cells (Figure 7A): all POLG^homo^ NSCs were fully able generate DA neurons whereas only 3/14 POLG^comp^ iPSCs clones gave established DA neurons (Figure 7A).

We tested this further in a three-dimensional (3D) system using the hanging drop technique to make 3D spheroids and iPSC-derived neuronal progenitors, astrocytes, and oligodendrocytes, as previously described (16). We found that both POLG spheroids were smaller than control lines, but POLG^comp^ spheroids were smaller in size and had less expression of the neural marker doublecortin (DCX) compared to POLG^homo^ spheroids (Figure 7B).

These data suggest that the more severe mitochondrial dysfunction found in compound heterozygous POLG variants results in a reduced neural differentiation potential compared to homozygous variants.

### Compound heterozygous POLG fibroblasts had similar mitochondrial function compared to homozygous POLG fibroblasts

We demonstrated above that mitochondrial function were more damaged in compound heterozygous neurons, and then we wondered whether this difference was manifested in parental fibroblasts. We then performed the same functional studies to examine mitochondrial-related features.

When comparing POLG^comp^ fibroblasts with POLG^homo^ fibroblasts, we observed similar cell morphology observed under phase contrast microscopy and mitochondrial morphology observed by MTG staining under confocal microscopy (Figure 7C). Flow cytometry analysis using double staining of TMRE and MTG showed similar mitochondrial mass (MTG), total MMP levels measured by TMRE and specific MMP level quantified by the ratio of total TMRE to MTG in the two patient fibroblasts (Figure 7D). ATP production measurements showed that ATP level, total and specific ROS levels (quantified by the ratio of total DCFDA per MTDR) in POLG^comp^ and POLG^homo^ fibroblasts were similar (Figure 7E).

We then compared changes in mtDNA copy number in fibroblasts from these two patients using the same methods described above. Flow cytometry analysis showed that both patients had similar total and specific levels of TFAM (Figure 7F), as well as for TOMM20 level, indicating the same mitochondrial mass (Figure 7F). QPCR also revealed similar mtDNA copy number in both mutant fibroblasts (Figure 7F). We then determined the protein levels of mitochondrial complexes and found no significant differences in total and specific levels of complexes I, III and IV between the two patient fibroblasts (Figure 7G).

These results suggest that compound heterozygous and homozygous POLG variants display similar mitochondrial function in fibroblasts.

## Discussion

Mutations in *POLG* give syndromes with variable clinical features and interestingly, different degrees of severity. While much has been learned about the cellular consequences of *POLG* mutations, establishing genotype-phenotype correlations have remained elusive. To address this question, we used reprogramming of patient-specific iPSCs to assess mutation-specific neuropathology. We generated iPSCs from from two patients, one homozygous for c.2243G>C (p.W748S) and the other compound heterozygous for c.2243G>C and c.1399G>A (p.A467T) mutations. Our earlier clinical studies showed that patients exhibit progressive neurological disease regardless of genotype, but major differences in survival could be observed depending on genotype, with compound heterozygotes having significantly shorter survival times than homozygous patients (4). In this present study, we found that compound heterozygous cells do indeed demonstrate a more severe defect in mitochondrial function than homozygous cells, findings supported by transcriptomic analysis showing more severe dysregulation of multiple metabolic pathways, but this only occurred in neural lineage cells, not in the original iPSCs or astrocytes.

The *POLG* mutants A467T and W748S affect the catalytic subunit through apparently distinct mechanisms. The W748S is located in the linker region, and it has been suggested that the W748S substitution would destabilize this subdomain and affect DNA polymerase activity (18). The W748S mutation causes low DNA polymerase activity, low DNA polymerase processivity, and defective DNA binding (19). In contrast, the A467T is located in the terminal thumb region of Pol γ in close contact with the dimeric auxiliary subunit (POLG2). It interacts with the template DNA and the helper subdomain and forms a major surface of the DNA binding channel (20). Functional studies have shown that the A467T mutant has both low polymerase activity and subunit interaction defects (21). That different mutations affect mtDNA replication defects in different ways raises the possibility that there are cumulative effects in compound heterozygotes resulting in a more severe phenotype than homozygotes. Further, since different tissues manifest different mtDNA defects in the presence of the same POLG mutations, e.g., neurons show mtDNA depletion while skeletal muscle shows multiple deletions, it is necessary to examine the question of genotype: phenotype correlation in specific tissues. Since it is not possible to do this in living patients, we turned to a tractable model system based on stem cell technology.

We have already shown that POLG iPSC derived NSCs replicate the cellular phenotypes found in postmortem neurons (5). In the current study, we show that, at the cellular level, the degree of mitochondrial DNA dysfunction in NSCs varies according to mutation: compound heterozygotes show a more pronounced mtDNA depletion and more impaired mitochondrial function, including MMP reduction, ATP production, and mitochondrial complex I level. Since complex I of the respiratory chain re-oxidizes reduced nicotinamide adenine dinucleotide (NADH) to NAD^+^, loss of complex I will have a major impact on metabolism generally. At the transcriptional level, we see this in our transcriptomic findings a clear difference between with compound heterozygotes and homozygous *POLG* mutations, with downregulation of mitochondrial function-related genes in compound heterozygotes, including genes in key mitochondrial functional modules involved in electron transport and energy production processes. Our findings suggest that more impaired mitochondrial function in heterogeneous NSCs may be reflected in core steps of mitochondrial energy production, and that these abnormalities ultimately lead to more severe ATP production defects compared to homozygotes, which further affects the differentiation capacity of these NSCs to generate committed neurons.

Transcriptomics revealed clear differences between compound heterozygotes and homozygous *POLG* NSCs, with downregulation of genes related to mitochondrial function in compound heterozygous NSCs, including genes in key mitochondrial functional modules involved in electron transport and energy production processes. Consistent with our functional studies, we found that genes encoding respiratory chain complex I, III, IV, and V proteins were downregulated, illustrating abnormalities in oxidative phosphorylation metabolism in heterozygous NSCs. Glycolysis, along with oxidative respiration, is the primary metabolic pathway employed by cells to transform biochemical energy from nutrients into ATP for cellular function. Pyruvate produced by glycolysis is transported to the mitochondria, where it enters the TCA cycle under and then is converted to lactate. Under hypoxic conditions or pathology, the rate of glycolysis increases to compensate for reduced oxidative respiration to meet cellular energy demands. We found that mitochondrial pyruvate and lactate metabolism-related pathways and genes were also significantly down-regulated in transcriptomics. These findings suggest that more impaired mitochondrial function in heterogeneous NSCs reflects not only core steps in mitochondrial energy production, especially oxidative phosphorylation, but also abnormalities in the pyruvate and lactate energy metabolism axes that ultimately lead to more severe ATP production deficiency in compound heterozygotes,

Mitochondrial dysfunction plays an important role in multiple feedback loops that induce cellular senescence *in vitro* (15) and *in vivo* (22). Cellular senescence is a complex stress response by which proliferating cells lose their ability to divide and enter cell cycle arrest. Growing evidence also suggests that senescent cells accumulate with age, a process that may contribute to age-related phenotypes and pathology (23). Several studies suggest that mitochondrial ROS are involved in this process; ROS can damage nuclear DNA, thereby activating the DNA damage response to induce senescence (15). On the other hand, metabolic decline can also lead to oxidative stress and DNA damage. We found that the more severe mitochondrial dysfunction and reduction of MMP in compound heterozygotes could lead to increased production of ROS and failure of NADH reoxidation, suggesting that both may be involved in the senescence response seen in these NSCs. While mitochondrial dysfunction drives and maintains cellular senescence (15, 24), senescence itself, particularly persistent DNA damage response signaling, also directly contributes to senescence-associated mitochondrial dysfunction (25). It is possible therefore, that both mechanisms are active in in POLG compound heterozygotes NSCs.

In addition to senescence, we found that another cellular consequence induced by the more severe mitochondrial damage in compound heterozygotes involved mitophagy. Mitophagy plays a role in maintaining mitochondrial health. Mitophagy can either specifically eliminate the damaged or dysfunctional mitochondria or clear all mitochondria during specialized developmental stages. The activation of mitophagy is observed in brain damage induced by cerebral ischemia (26). Similar, our finding on the upregulation of BNIP3 and LC3B protein levels and occurrence of mitochondrial autophagosomes in NSCs with compound heterozygous POLG mutations suggested active degradation through mitophagy. More importantly, given the mitophagy is an essential mitochondrial quality control mechanism implicated in senescence and it can also be induced by mitochondrial stress (27). Our finding of the more mitochondrial damage in NSCs for compound heterozygote than homozygote may lead to the cell fate of both mitophagy activation and senescence and later will even enhance the mitochondrial stress. Studies in stroke revealed that BNIP3 exclusively activates excessive mitophagy leading to cell death (28). Therefore, the activation of mitophagy at the NSC stage may lead to a decrease in the efficiency of DA neuron differentiation in POLG compound heterozygous cells.

In the present study, we found that patient fibroblasts and iPSCs in compound heterozygotes appeared to be able to maintain their MMP, mitochondrial mass, and mtDNA replication at similar levels as homozygotes (Figure S5). They also appear to be able to modulate their redox homeostasis. Interestingly, when NSCs were differentiated into astrocytes, mitochondrial function appeared relatively similar in compound heterozygotes and homozygotes, even though these cells were derived from NSCs with clear differences in the severity of the mitochondrial defect. This is similar to what we see in fibroblasts and suggests that cells such as these, that are highly dependent on glycolysis for energy generation, are in some way protected from the most damaging consequences.

In conclusion, our study shows that it is possible to construct a genotype-phenotype correlation to explain why patients with compound heterozygotes mutations have a poorer prognosis than those with homozygous *POLG* mutations. In compound heterozygotes, mitochondrial function is more impaired than homozygotes and the effects on other metabolic pathways particularly energy metabolism, are more profound. We find also that there is also greater activation of cellular senescence and pathways required for clearing damaged mitochondria (Figure 7H). That this is restricted to neural lineage cells, and for example not seen in astrocytes, is also important. Our observations from postmortem studies showed that a major astrocyte response to the neuronal damage and necrosis suggesting that this cell type remains functional. Whether this difference can be explained by two mutations affecting polymerase function in different ways remains unclear. We did, however, find lower mtDNA copy number in heterozygous cells suggesting that this possibility remains.

## Material and methods

### Ethical permission

The project was approved by the Western Norway Committee for Ethics in Health Research (REK nr. 2012/919). Tissues were acquired with written informed consent from all patients.

### Generation of iPSCs, NSCs, DA neurons and astrocytes

Skin biopsies were collected from POLG patients and fibroblasts were isolated. Ethical approval was granted from Norwegian Research Ethics Committee (2012/919). Fibroblasts were cultured in medium consisted of DMEM/F12, GlutaMAX™ (Thermo Fisher Scientific, # 35050061), and 10% (v/v) Fetal Bovin Serum (FBS, Sigma-Aldrich, # 12103C). Fibroblasts were reprogrammed to iPSCs as previously described (5). iPSCs were grown under feeder-free condition using E8 medium (Invitrogen, # A1517001) on 6 well Geltrex-coated plates. The generation of NSCs and DA neurons were described previously (5). Astrocytes were differentiated from NSCs according to the protocol described previously (16).

### Immunocytochemistry and immunofluorescence staining

Live cells grown on coverslips were fixed with 4% (v/v) paraformaldehyde (PFA) for 10 min. Then the cells were blocked using blocking buffer composed of 1X PBS, 10% (v/v) normal goat serum (Sigma-Aldrich, # G9023) and 0.3% (v/v) Triton X-100 (Sigma-Aldrich, # X100-100ML) for 1 h. The cells were then incubated with primary antibody at 4°C for overnight. The following antibodies were used: rabbit anti-NANOG (Abcam, # ab80892), rabbit anti-OCT4 (Abcam, # 19857), rabbit anti-SOX2 (Abcam, # ab97959), rabbit anti-PAX6 (Abcam, # ab5790), rabbit anti-SOX2 and mouse anti-NESTIN (Santa Cruz Biotechnology, # sc23927), chicken anti-GFAP (Abcam # ab4674), rabbit anti-S100ß (Abcam, # ab196442), rabbit anti-EAAT-1 (Abcam, # ab416, 1:200) and mouse anti-Glutamine synthetase (Abcam, # ab64613, 1:200). After three washing with PBS, cells were further incubated with secondary antibody (1:800 in blocking buffer) for 1 h at RT. The secondary antibodies included Alexa Flour® goat anti-mouse 594 (Thermo Fisher Scientific Scientific, # A11005, 1:800) and Alexa Flour® goat anti-rabbit 488 (Thermo Fisher Scientific Scientific, # A11008, 1:800). The coverslips were mounted using Prolong Diamond Antifade Mountant with DAPI (Thermo Fisher Scientific, # P36962).

### Intracellular ATP production

Intracellular ATP production was detected using the Luminescent ATP Detection Assay Kit (Abcam, cat.number: ab113849). Cells were grown to 90% confluency in 96 well plate (Life Sciences, # 3601). ATP measurements were conducted based on manufacturer’s protocol. Cells were lysed. Luciferase enzyme and luciferin were added. Emitted light corresponding to the number of ATPs was measured in a Victor® XLight Multimode Plate Reader (PerkinElmer). For each sample, 3-6 replicates were measured. In order to normalize ATP production by cell numbers, cells grown in the same 96 well plates were stained using Janus Green cell normalization stain kit (Abcam, # ab111622). OD value at 595 nm was measured by Labsystems Multiskan Bichromatic plate reader (Titertek Instruments, USA).

### ROS production

ROS production was measured by flow cytometry. Intracellular ROS production in relation to mitochondrial mass was quantified using double staining of 30 μM dichlorofluorescin diacetate (DCFDA) (Abcam, # ab11385) and 150 nM MitoTracker Deep Red (MTDR) (Invitrogen, # M22426). Mitochondrial ROS production in relation to mitochondrial mass was detected using double staining of 10 μM MitoSOX red mitochondrial superoxide indicator (Thermo Fisher Scientific, # M36008) and 150 nM MTG. The stained cells were detached using TrypLE Express enzyme and neutralized using culture media with 10% FBS, before being analyzed on a FACS BD Accuri™ C6 flow cytometer. More than 4,000 cells were recorded for each sample. results were analyzed using BD Accuri C6 Software.

### NADH and NAD^+^ measurement

NADH and NAD^+^ were measured using liquid chromatography mass spectrometry (LC-MS). After washing with PBS, cells were incubated with ice-cold 80% methanol for 20 min at 4°C and were stored at −80°C overnight. The next day, samples were thawed on a rotating wheel at 4°C and subsequently centrifuged at 16 000 g for 20 min at 4°C. The supernatant was added to 1 volume of acetonitrile and were stored at −80°C until analysis. The cell pellet was dried and reconstituted in lysis buffer (20 mM Tris-HCl (pH 7.4), 2% SDS, 1 mM EDTA, 150 mM NaCl) for protein determination (BCA assay). Metabolite separation was achieved by using a ZIC-pHILC column (150 x 4.6 mm, 5 μm; Merck) and Dionex UltiMate 3000 liquid chromatography system (Thermo Fisher Scientific). The column temperature was kept at 30°C. The mobile phase consisted of 10 mM ammonium acetate pH 6.8 (Buffer A) and acetonitrile (Buffer B). The flow rate was kept at 400 μL/min, and the gradient was set at 0 min 20% Buffer B, 15-20 min 60% Buffer B, 35 min 20% Buffer B. Ionization was achieved by heated electrospray ionization (HESI-II) probe (Thermo Fisher Scientific) using the positive ion polarity mode, with a spray voltage of 3.5 kV. The sheath gas flow rate was 48 units. The auxiliary gas flow rate was 11 units. The sweep gas flow rate was 2 units. The capillary temperature was 256°C. The auxiliary gas heater temperature was 413°C. The stacked-ring ion guide (S-lens) radio frequency level was at 90 units. Mass spectra were recorded by QExactive mass spectrometer (Thermo Fisher Scientific). Data analysis was performed using Thermo Xcalibur Qual Browser (Thermo Fisher Scientific). Standard curves generated for NAD^+^ and NADH were used as quantification references.

### Measurement of mtDNA copy number

MtDNA copy number was quantified using both flow cytometry and qPCR. For flow cytometry, cells were double stained with TFAM and TOMM20. Cells were detached with TrypLETM Express enzyme and fixed with 1.6% (v/v) PFA (VWR, # 100503 917) for 10 minutes at RT. Thereafter, cells were permeabilized with ice-cold 90% methanol for 20 min at −20°C, and blocked-in blocking buffer containing PBS, 0.3M glycine, 5% goat serum and 1% bovine serum albumin (BSA). Cells were incubated with anti-TFAM antibody conjugated to Alexa Fluor® 488 (Abcam, # ab198308) at 1:400 dilutions and anti-TOMM20 antibody conjugated to Alexa Fluor® 488 (Santa Cruz Biotechnology, # sc 17764 AF488) at 1:400 dilutions. The cells were then analyzed on BD AccuriTM C6 flow cytometer and data analysis was performed using BD Accuri C6 Software. The mtDNA quantities relative to mitochondria mass were represented by TFAM/TOMM20 ratio.

For qPCR, DNA was extracted using a QIAGEN DNeasy Blood and Tissue Kit (QIAGEN, # 69504) according to the manufacturer’s protocol. MtDNA was quantified as previously described (5, 6).

### Measurement of complex I, II, and IV expression

Protein levels of mitochondrial complexes I, II, and IV were accessed using flow cytometry. Cells were detached with TrypLE enzyme and were fixed with 1.6% (v/v) PFA (VWR, # 100503 917) for 10 minutes at RT, before permeabilized using ice-cold 90% methanol at −20°C for 20 min. The cells were blocked using the blocking buffer mentioned above. Cells were stained with primary antibodies at dilutions 1:1000. The primary antibodies include anti-NDB10 (Abcam, # ab196019), anti-SDHA (Abcam, # ab14715) and anti-COX IV (Abcam, # ab14744). The secondary antibodies were subsequently incubated at dilutions 1:400. The cells were analyzed on BD Accuri™ C6 flow cytometer and data analysis was performed using BD Accuri C6 Software. At least 40,000 events were recorded for each sample, doublets or dead cells were excluded.

### MMP and mitochondrial mass measurement

MMP relative to mitochondrial mass was measured using flow cytometry. Cells were duel stained with 100 nM TMRE (Abcam, # ab113852) and 150 nM MTG (Invitrogen, # M7514) for 45 min at 37°C. Cells treated with 100 μM FCCP (Abcam, # ab120081) was used as negative control. After washing with PBS, cells were detached with TrypLE™ enzyme and neutralized with culture media plus 10% FBS. The cells were then analyzed on a FACS BD Accuri™ C6 flow cytometer (BD Biosciences, San Jose, CA, USA). The data analysis was performed using BD Accuri C6 Software. At least 40,000 events were recorded for each sample, doublets or dead cells were excluded.

### Bulk RNA sequencing analysis

Total RNA was isolated from NSCs and iPSCs using RNeasy Mini Kit (QIAGEN). RNA quality and concentration were checked using Bioanalyzer and Qubit™. RNA sequencing libraries were established by the PolyA enrichment method. For iPSCs and NSCs, sequencing was performed by HudsonAlpha. FASTQ files were trimmed using Trimmomatic version 0.39 to remove potential Illumina adapters and low-quality bases with the following parameters: ILLUMINACLIP: truseq. fa: 2:30:10 LEADING:3 TRAILING:3 SLIDINGWINDOW: 4:15 (29). FASTQ files were assessed using fastQC version 0.11.8 prior and following trimming. We used Salmon version 1.0.0 (30) to quantify the abundance at the transcript level with the fragment-level GC bias correction option (*gcBias*flag) and the appropriate option for the library type (*l*flag set to A) against the GENCODE release 32 of the human transcriptome (GRCh38.p13). Transcript-level quantification was imported into R and collapsed onto gene-level quantification using the tximport R package version 1.8.0 (31) according to the gene definitions provided by the same GENCODE release. We filtered out genes in non-canonical chromosomes and scaffolds, and transcripts encoded by the mitochondrial genome. Genes were filtered out if their level of expression was below 10 reads in more than 75% of the samples based on CPM (counts per million) (32), which resulted in a total of 20,565 genes for downstream analyses. Transcripts counts were aggregated into gene counts. Sample correlations were calculated with the information of DV200, library size, cell type and mutation, and sample outliers were excluded (Figure S2). Differential expression analysis was conducted using DEseq2 (33). Multiple comparisons were adjusted by using the false discovery rate method. Adjusted P value (q value) <0.05 was considered as statistical significance. KEGG pathway and KEGG module enrichment analysis were conducted using Clusterprofiler (34). Human MitoCarta 2.0 was used as a reference database of collections of genes with mitochondrial related function and localization (13). For astrocytes, RNA sequencing was performed by Beijing Genomics Institute (BGI, Shenzhen, China). Library preparation was conducted at BGI following the guide of the standard protocol. The sequencing was performed at BGI-Shenzhen using BGISEQ-500. Low-quality reads and adapter contaminated reads were removed using SOAPnuke software (35). The processed fastq files were mapped to the human genome using HISAT (36). The genome version was GRCh38, with annotations from Bowtie2 (37). Then the expression level of the gene was calculated by RSEM (v1.2.12) (38). Differential expression was conducted using the DEseq2 package.

### Statistical analysis

All data are expressed as mean ± standard error of the mean (SEM). Statistical analysis was performed using GraphPad Prism 7.0 (GraphPad Software). For a data set that meets the normal distribution, two-side student’s t test was applied, otherwise the Mann-Whitney U test was applied. *P* values less than 0.05 was considered statistical significance.

## Data Availability

The datasets generated and analyzed during the study are included with the Supplemental Information. The RNA sequencing read count data can be accessed in NCBI Gene Expression Omnibus (GEO) data deposit system with an accession number GSE207007. All other data are available from the corresponding author upon request.

## Conflict of Interest

All authors declare that the research was conducted in the absence of any commercial or financial relationships that could be construed as a potential conflict of interest.

## Funding

This work was supported by the following funding: K.L was partly supported by University of Bergen Meltzers Høyskolefonds (project number:103517133) and Gerda Meyer Nyquist Legat (project number: 103816102). L.A. B was supported the Norwegian Research Council (project number: 229652), Rakel og Otto Kr.Bruuns legat. G.J.S was partly supported by the Norwegian Research Council through its Centres of Excellence funding scheme (project number: 262613).

## Author’s contributions

K.L contribute to the conceptualization; K.L, Y. H and C.K.K contribute to the methodology; K.L, Y. H, C.K.K, A.C, G.N and L.E.H contribute to the investigation; K.L and Y. H contribute to the writing original draft; all authors contribute to writing review and editing; K.L, L.A.B and G.J.S contribute to the funding acquisition; K.L, G.J.S, L.A.B and M. Z contribute to the resources; K.L contributes to the supervision. All authors agree to the authorships.

## Acknowledgements

We thank members of the Molecular Imaging Centre, Flow Cytometry Core Facility and Genomics Core Facility for their expertise and assistance in confocal imaging and flow cytometry data recording, generating the DNA sequencing data.

---

**Figure S1.**
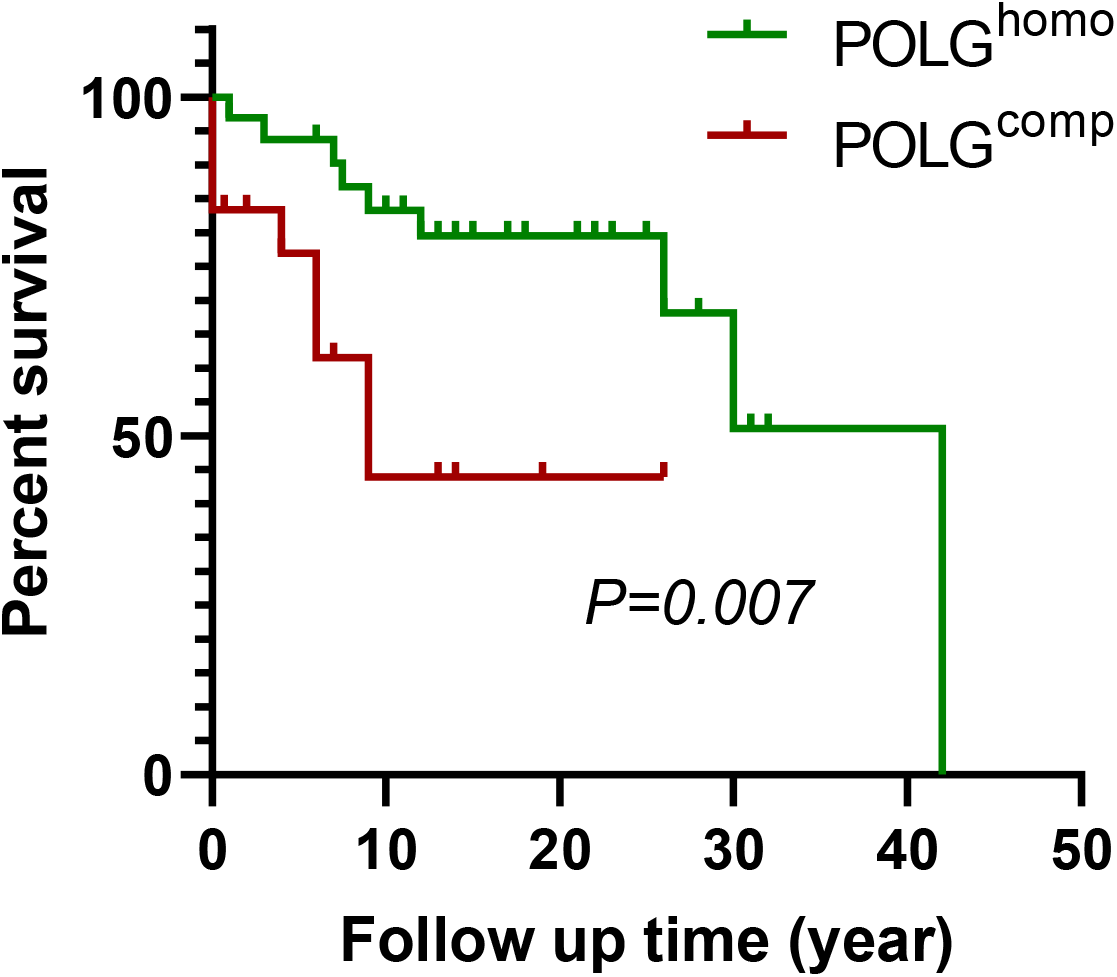
Survival curves of POLG patients carrying POLG^homo^ and POLG^comp^ in collected studies (P=0.007).

**Figure S2.**
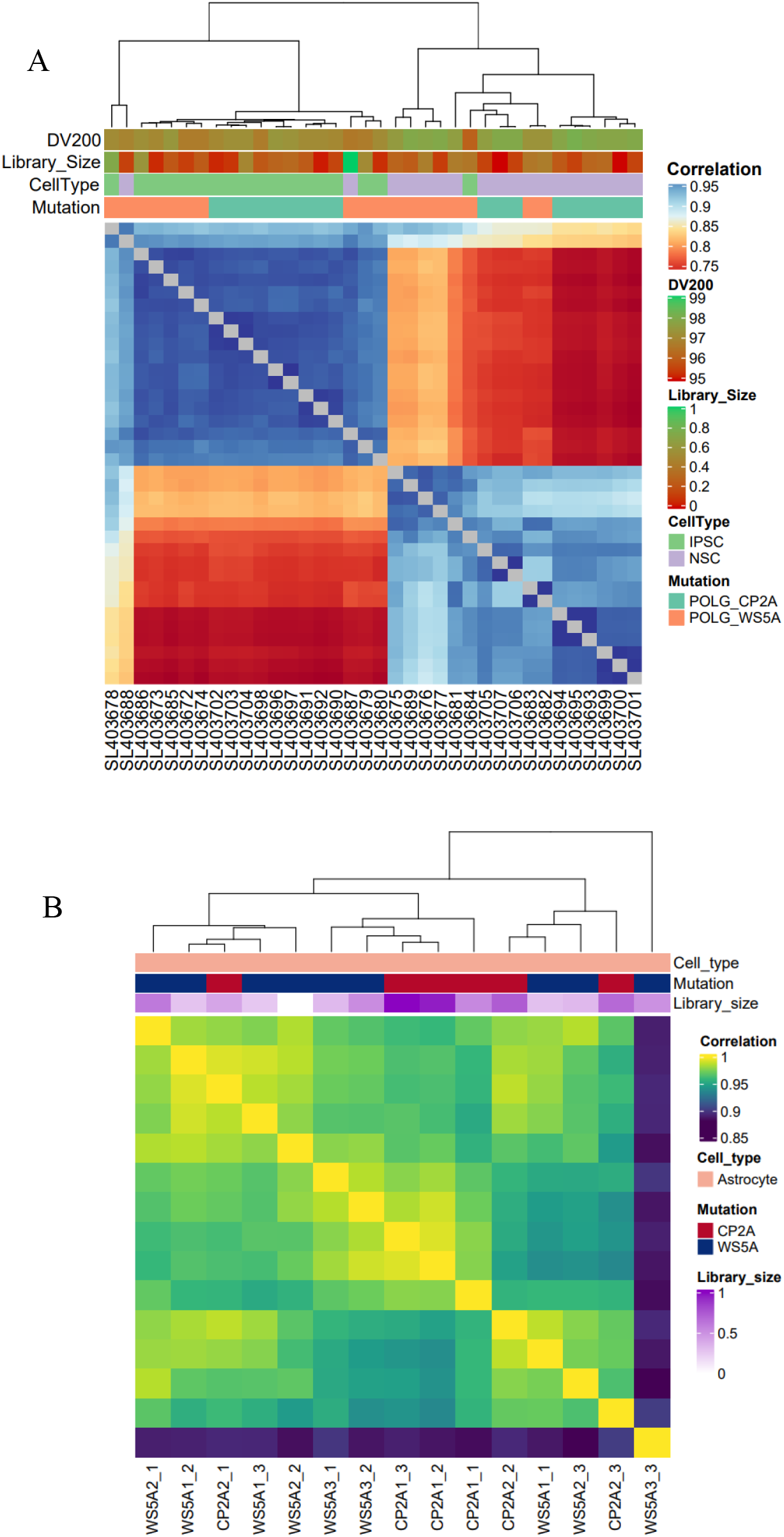
Sample correlations in RNA-sequencing analysis in POLG^homo^ and POLG^comp^ iPSCs, NSCs (A) and astrocytes (B).

**Figure S3.**
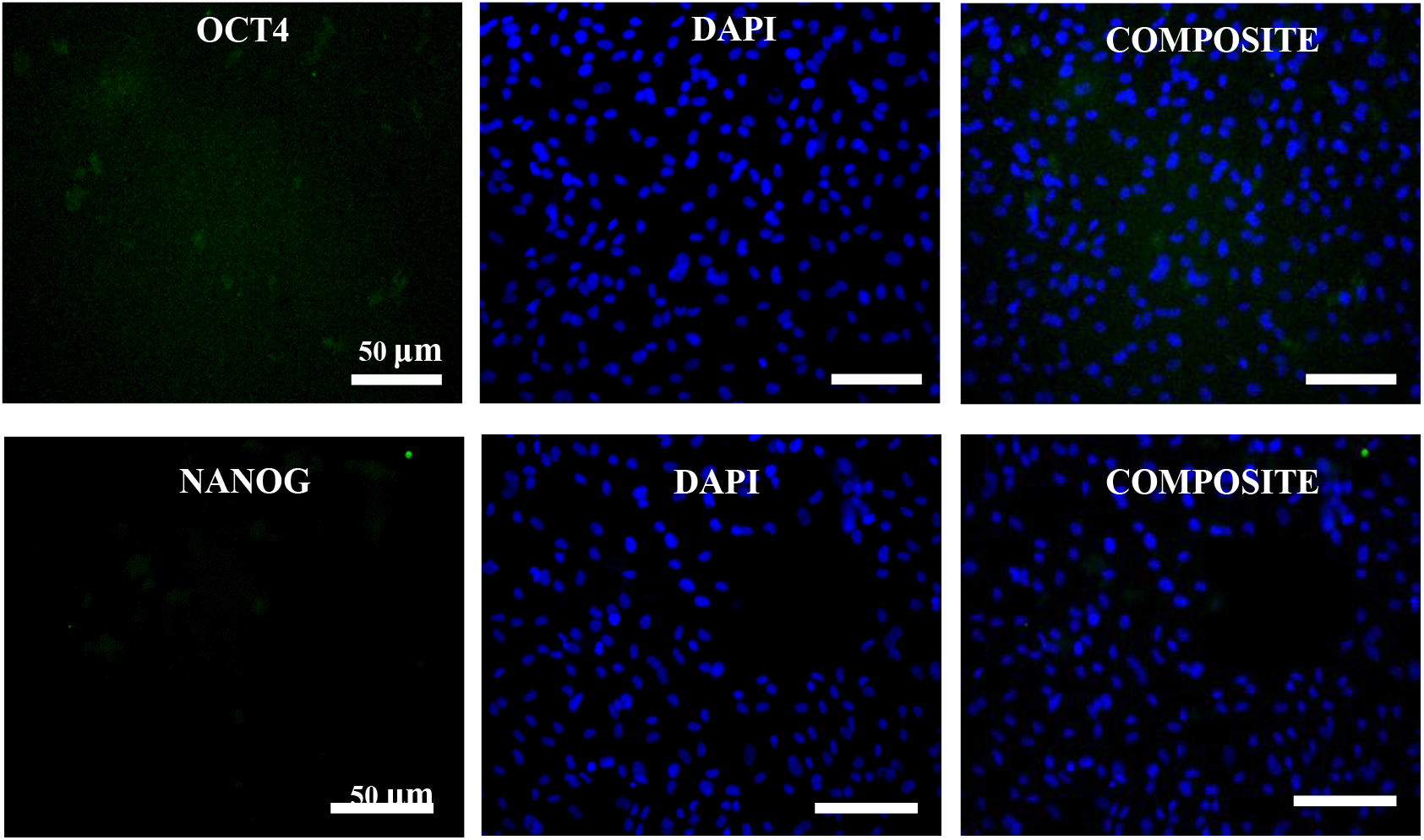
Representative images of pluripotent stem cell marker OCT4 and IiPSC-derived NCSs.

**Figure S4.**
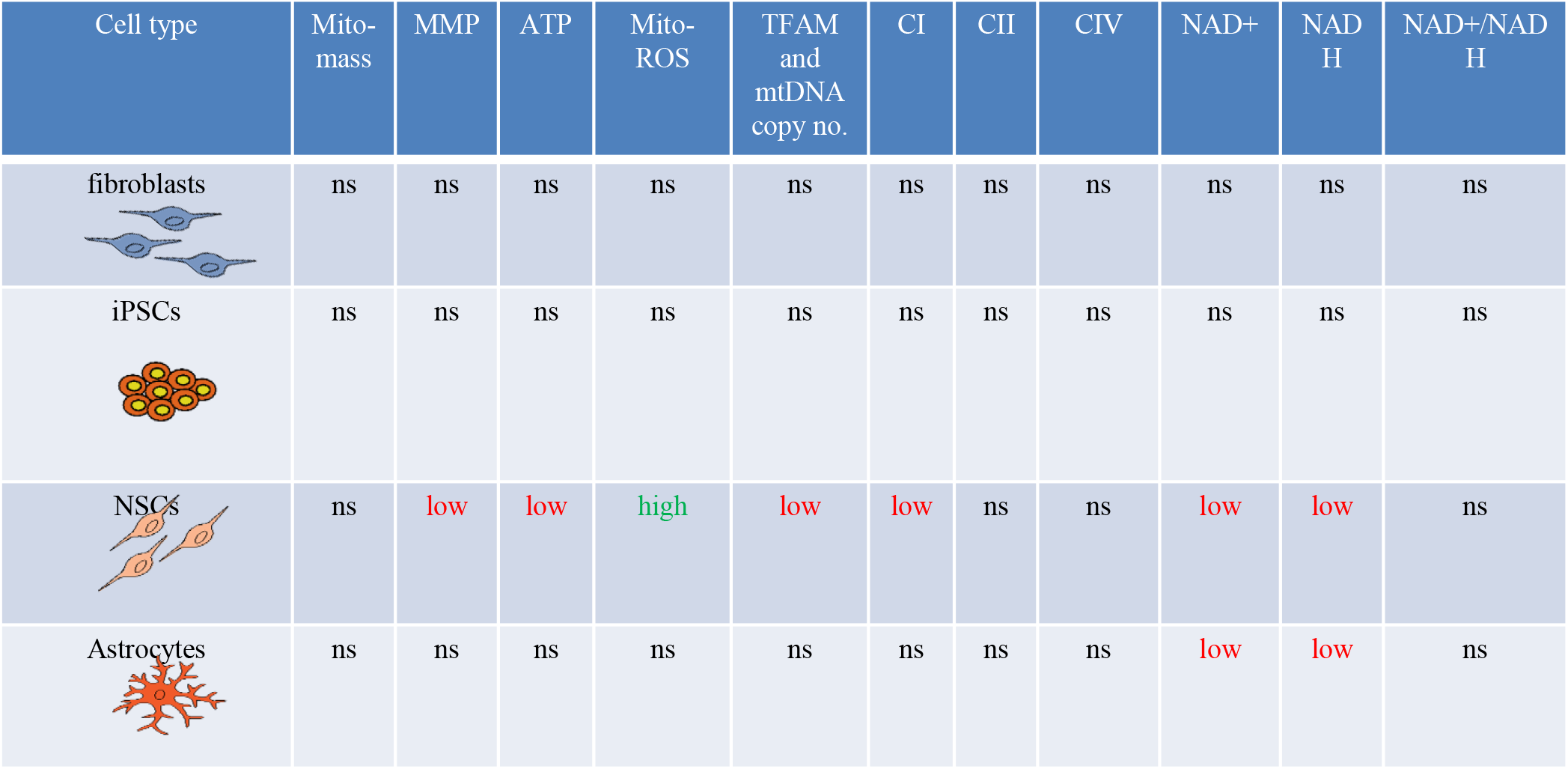
Comparation of the changes in mitochondrial-related measures in different cell types when comparing cells compozygous versus homozygous for POLG compounds. ns: no significant change.

**Table S1.** Included studies for systemic evaluation and comparison of POLG^comp^ and POLG^homo^ phenotypes.

| **Study** | **Genotype** | **Patient number** | **Country** |
| --- | --- | --- | --- |
| Tzoulis et al, 2006 (7) | A467T/W748S | 7 | Norway |
|  | W748S/W748S | 13 |  |
| Hakonen et al, 2005 (8) | W748S/W748S | 19 | Finland |
| Lax et al, 2012 (9) | A467T/W748S | 2 | United Kingdom |
| Janssen et al, 2016 (10) | W748S/W748S | 2 | Belgium |
|  | A467T/W748S | 3 |  |
| Kollberg et al, 2006 (11) | A467T/W748S | 2 | Sweden |

**Table S2.**
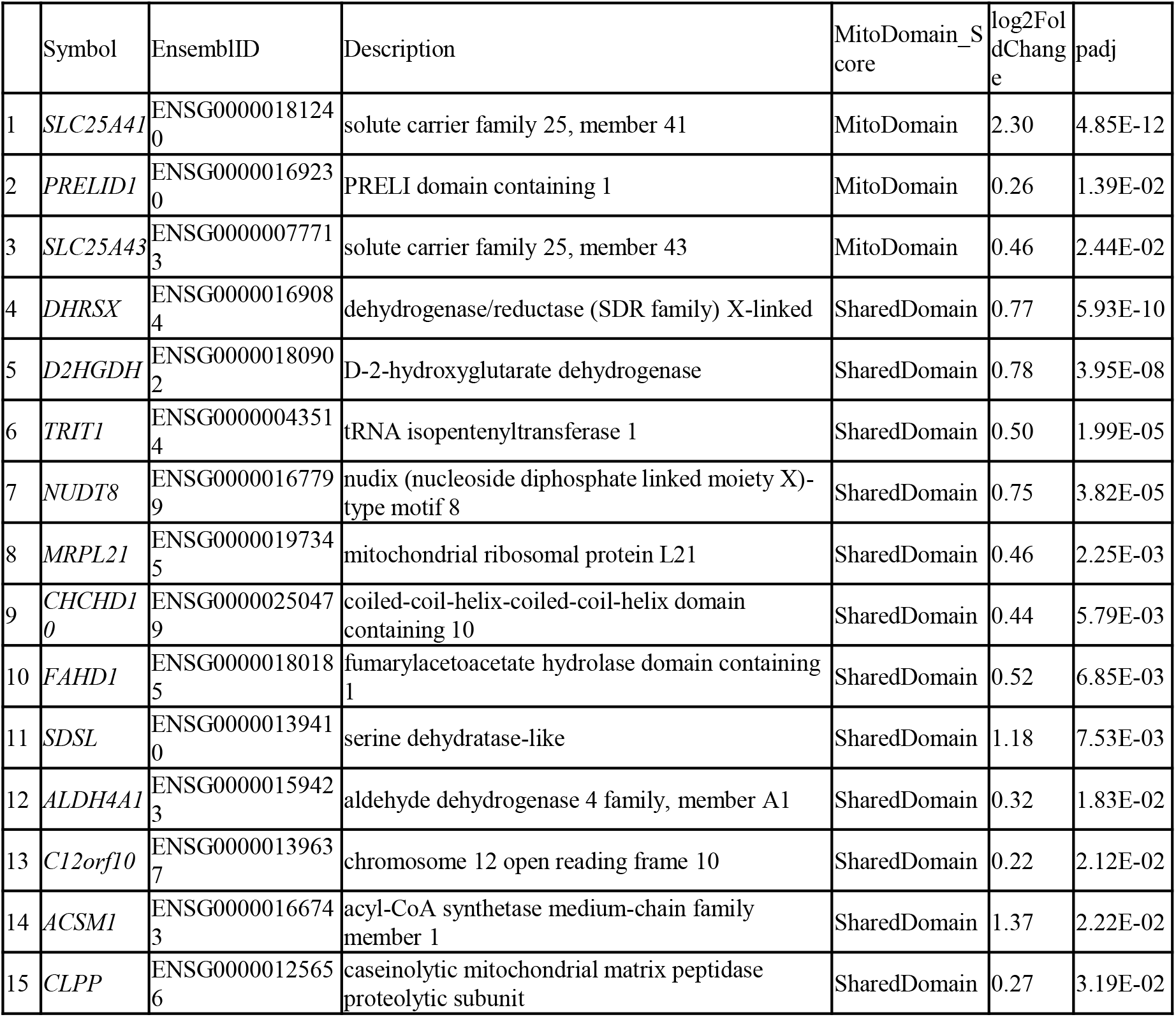
Top 15 upregulated mitochondrial genes in genes in POLG^comp^ iPSCs compared with POLG^homo^ iPSCs.

**Table S3.** Top 15 downregulated mitochondrial genes in POLG^comp^ compared with POLG^homo^ iPSCs.

|  | Symbol | EnsemblID | Description | MitoDomain_Score | log2FoldChange | padj |
| --- | --- | --- | --- | --- | --- | --- |
| 1 | *CPT1C* | ENSG00000169169 | carnitine palmitoyltransferase 1C | SharedDomain | -0.60 | 8.12E-12 |
| 2 | *PABPC5* | ENSG00000174740 | poly(A) binding protein, cytoplasmic 5 | SharedDomain | -1.55 | 6.33E-10 |
| 3 | *PITRM1* | ENSG00000107959 | pitrilysin metallopeptidase 1 | MitoDomain | -0.60 | 4.59E-06 |
| 4 | *DHRS1* | ENSG00000157379 | dehydrogenase/reductase (SDR family) member 1 | SharedDomain | -0.64 | 1.09E-05 |
| 5 | *PDK3* | ENSG00000067992 | pyruvate dehydrogenase kinase, isozyme 3 | MitoDomain | -0.95 | 2.04E-05 |
| 6 | *CDC25C* | ENSG00000158402 | cell division cycle 25C | SharedDomain | -0.46 | 3.51E-05 |
| 7 | *LYRM5* | ENSG00000205707 | LYR motif containing 5 | MitoDomain | -0.58 | 6.00E-05 |
| 8 | *NDUFS5* | ENSG00000168653 | NADH dehydrogenase (ubiquinone) Fe-S protein 5, 15kDa (NADH-coenzyme Q reductase) | MitoDomain | -0.56 | 1.36E-04 |
| 9 | *SURF1* | ENSG00000148290 | surfeit 1 | MitoDomain | -0.42 | 6.73E-03 |
| 10 | *COA5* | ENSG00000183513 | cytochrome c oxidase assembly factor 5 | MitoDomain | -0.39 | 7.76E-03 |
| 11 | *LYRM4* | ENSG00000214113 | LYR motif containing 4 | MitoDomain | -0.33 | 7.87E-03 |
| 12 | *MRPS33* | ENSG00000090263 | mitochondrial ribosomal protein S33 | MitoDomain | -0.32 | 2.08E-02 |
| 13 | *CCDC90B* | ENSG00000137500 | coiled-coil domain containing 90B | MitoDomain | -0.37 | 3.35E-02 |
| 14 | *SLC25A4* | ENSG00000151729 | solute carrier family 25 (mitochondrial carrier; adenine nucleotide translocator), member 4 | MitoDomain | -0.28 | 4.43E-02 |
| 15 | *SLC25A14* | ENSG00000102078 | solute carrier family 25 (mitochondrial carrier, brain), member 14 | MitoDomain | -0.68 | 4.83E-02 |

Note: only mitoDomain genes were listed. E-: X 10

**Table S4.** Top 15 upregulated mitochondrial genes in POLG^comp^ compared with POLG^homo^ NSCs.

|  | Symbol | EnsemblID | Description | MitoDomain_Score | log2Fold Change | padj |
| --- | --- | --- | --- | --- | --- | --- |
| 1 | *GLS* | ENSG00000115419 | glutaminase | MitoDomain | 0.87 | 2.39E-13 |
| 2 | *SLC25A43* | ENSG00000077713 | solute carrier family 25, member 43 | MitoDomain | 0.93 | 1.10E-12 |
| 3 | *SLC25A45* | ENSG00000162241 | solute carrier family 25, member 45 | MitoDomain | 1.82 | 1.21E-09 |
| 4 | *TOMM6* | ENSG00000124593 | translocase of outer mitochondrial membrane 6 homolog (yeast) | MitoDomain | 1.31 | 1.12E-07 |
| 5 | *FIS1* | ENSG00000214253 | fission 1 (mitochondrial outer membrane) homolog (S. cerevisiae) | MitoDomain | 0.32 | 1.30E-06 |
| 6 | *MUT* | ENSG00000146085 | methylmalonyl CoA mutase | MitoDomain | 0.52 | 1.83E-06 |
| 7 | *SLC25A16* | ENSG00000122912 | solute carrier family 25 (mitochondrial carrier; Graves disease autoantigen), member 16 | MitoDomain | 0.40 | 3.74E-06 |
| 8 | *ACAD11* | ENSG00000240303 | acyl-CoA dehydrogenase family, member 11 | MitoDomain | 0.68 | 1.04E-04 |
| 9 | *MUL1* | ENSG00000090432 | mitochondrial E3 ubiquitin protein ligase 1 | MitoDomain | 0.48 | 1.17E-04 |
| 10 | *BCKDHB* | ENSG00000083123 | branched chain keto acid dehydrogenase E1, beta polypeptide | MitoDomain | 0.54 | 1.32E-04 |
| 11 | *BNIP3* | ENSG00000176171 | BCL2/adenovirus E1B 19kDa interacting protein 3 | MitoDomain | 1.86 | 3.17E-04 |
| 12 | *SLC25A46* | ENSG00000164209 | solute carrier family 25, member 46 | MitoDomain | 0.20 | 4.03E-04 |
| 13 | *PDK3* | ENSG00000067992 | pyruvate dehydrogenase kinase, isozyme 3 | MitoDomain | 1.16 | 7.89E-04 |
| 14 | *GSTK1* | ENSG00000197448 | glutathione S-transferase kappa 1 | MitoDomain | 0.40 | 9.77E-04 |
| 15 | *GRPEL2* | ENSG00000164284 | GrpE-like 2, mitochondrial (E. coli) | MitoDomain | 0.33 | 1.32E-03 |

Note: only mitoDomain genes were listed. E-: X 10-

**Table S5.** Top 15 downregulated mitochondrial genes in POLG^comp^ compared with POLG^homo^ NSCs.

|  | Symbol | EnsemblID | Description | MitoDomain_Score | log2Fold Change | padj |
| --- | --- | --- | --- | --- | --- | --- |
| 1 | *NDUFS5* | ENSG00000168653 | NADH dehydrogenase (ubiquinone) Fe-S protein 5, 15kDa (NADH-coenzyme Q reductase) | MitoDomain | 1.22 | 8.53E-20 |
| 2 | *ATP5O* | ENSG00000241837 | ATP synthase, H<sup>+</sup> transporting, mitochondrial F1 complex, O subunit | MitoDomain | 0.63 | 5.56E-16 |
| 3 | *COX6B1* | ENSG00000126267 | cytochrome c oxidase subunit VIb polypeptide 1 (ubiquitous) | MitoDomain | 0.37 | 3.46E-15 |
| 4 | *PTCD1* | ENSG00000106246 | pentatricopeptide repeat domain 1 | MitoDomain | 0.69 | 5.99E-14 |
| 5 | *ATP5J2-PTCD1* | ENSG00000106246 | ATP5J2-PTCD1 readthrough | MitoDomain | 0.69 | 5.99E-14 |
| 6 | *UQCRH* | ENSG00000173660 | ubiquinol-cytochrome c reductase hinge protein | MitoDomain | 0.46 | 9.78E-14 |
| 7 | *MRPL51* | ENSG00000111639 | mitochondrial ribosomal protein L51 | MitoDomain | 0.65 | 2.44E-10 |
| 8 | *SLC25A17* | ENSG00000100372 | solute carrier family 25 (mitochondrial carrier; peroxisomal membrane protein, 34kDa), member 17 | MitoDomain | 0.56 | 3.31E-09 |
| 9 | *PTCD3* | ENSG00000132300 | pentatricopeptide repeat domain 3 | MitoDomain | 0.50 | 2.71E-08 |
| 10 | *DIABLO* | ENSG00000184047 | diablo, IAP-binding mitochondrial protein | MitoDomain | 0.34 | 3.40E-08 |
| 11 | *ATP5L* | ENSG00000167283 | ATP synthase, H<sup>+</sup> transporting, mitochondrial Fo complex, subunit G | MitoDomain | 0.44 | 1.28E-07 |
| 12 | *PDHA1* | ENSG00000131828 | pyruvate dehydrogenase (lipoamide) alpha 1 | MitoDomain | 0.52 | 1.41E-07 |
| 13 | *ATPAF1* | ENSG00000123472 | ATP synthase mitochondrial F1 complex assembly factor 1 | MitoDomain | 0.39 | 2.01E-07 |
| 14 | *MRPL50* | ENSG00000136897 | mitochondrial ribosomal protein L50 | MitoDomain | 0.70 | 3.21E-07 |
| 15 | *MRPL53* | ENSG00000204822 | mitochondrial ribosomal protein L53 | MitoDomain | 0.40 | 3.51E-07 |

Note: only mitoDomain genes were listed. E-: X 10-

**Table S6.** Upregulated mitochondrial genes in POLG^comp^ compared with POLG^homo^ astrocytes.

| Symbol | EnsemblID | Description | MitoDomain_Score | log2FoldChange | padj |
| --- | --- | --- | --- | --- | --- |
| *HSPB7* | ENSG00000173641 | heat shock 27kDa protein family, member 7 | SharedDomain | 6.34 | 3.99E-3 |

Note: There were no DE mitochondrial related genes found to be downregulated in compound compozygotes A467T/A467T compared with W748S homozygote. E-: X 10-

## Notes

### Competing Interest Statement

The authors have declared no competing interest.

